# Strong spatial population structure shapes the temporal coevolutionary dynamics of costly female preference and male display

**DOI:** 10.1101/2021.09.29.462412

**Authors:** Maximilian Tschol, Jane M. Reid, Greta Bocedi

## Abstract

Female mating preferences for exaggerated male display traits are commonplace. Yet, comprehensive understanding of the evolution and persistence of costly female preference through indirect (Fisherian) selection in finite populations requires some explanation for the persistence of additive genetic variance (V_a_) underlying sexual traits, given that directional preference is expected to deplete V_a_ in display and hence halt preference evolution. However, the degree to which V_a_, and hence preference-display coevolution, may be prolonged by spatially variable sexual selection arising solely from limited gene flow and genetic drift within spatially structured populations has not been examined. Our genetically and spatially explicit model shows that spatial population structure arising in an ecologically homogeneous environment can facilitate evolution and long-term persistence of costly preference given small subpopulations and low dispersal probabilities. Here, genetic drift initially creates spatial variation in female preference, leading to persistence of V_a_ in display through “migration-bias” of genotypes maladapted to emerging local sexual selection, thus fuelling coevolution of costly preference and display. However, costs of sexual selection increased the probability of subpopulation extinction, limiting persistence of high preference-display genotypes. Understanding long-term dynamics of sexual selection systems therefore requires joint consideration of coevolution of sexual traits and metapopulation dynamics.

## Introduction

Directional female mating preferences for elaborate and costly male display traits are widespread in nature (Andersson 1994; Wiens and Tuschhoff 2020). Explaining the evolution and long-term persistence of such female preferences and associated male displays has long challenged evolutionary biologists (Darwin 1871), especially when preferences are apparently costly with no clear direct fitness benefits (Pomiankowski 1987; Andersson 1994; Jennions and Petrie 1997; Friberg and Arnqvist 2003; Wong and Candolin 2005). Benefits of female preferences are then hypothesised to be indirect, stemming from genetic covariance between preference and male display that arises given initial additive genetic variance (V_a_) in both traits and resulting non-random mating (i.e., “Fisherian sexual selection”; Fisher 1915; 1930; Lande 1981; Kirkpatrick 1982). Given such additive genetic (co)variances, male displays might be expected to evolve such that survival costs (i.e., viability or natural selection; Lande 1981) are balanced by reproductive benefits through sexual selection.

Indeed, diverse models show that female preference can evolve via such indirect selection, depending on the strength of direct selection against preference and the shape of the variance-covariance matrix (G-matrix) for female preference and male display, and viability (Lande 1981; Kirkpatrick 1982; Kokko et al. 2002; Kokko et al. 2006; Henshaw and Jones 2020). However, one persistent challenge is that directional sexual selection is expected to quickly deplete V_a_ in male display traits and associated fitness (the “Lek paradox”), eliminating indirect selection for costly female preference (Borgia 1979; Kirkpatrick and Ryan 1991). Identifying mechanisms that help maintain V_a_ over long biological timeframes therefore remains key to explaining the continued expression of costly directional mating preference and resulting extraordinary phenotypic diversity (Mead and Arnold 2004; Tomkins et al. 2004; Kokko et al. 2006; Kotiaho et al. 2008; Radwan 2008; Prokuda and Roff 2014; Radwan et al. 2016; Lindsay et al. 2019).

Diverse mechanisms that could maintain V_a_ in male display have been proposed and tested (Radwan 2008; Bonilla et al. 2016; Radwan et al. 2016; Dugand et al. 2019). For example, V_a_ may be maintained by mutation-selection balance, whereby decreases due to sexual selection are compensated by sufficient frequency of new deleterious mutations (Iwasa et al. 1991; Pomiankowski et al. 1991), which could occur given a large enough mutational target (Pomiankowski and Møller 1995; Rowe and Houle 1996; Houle and Kondrashov 2002). Balancing selection may also maintain V_a_ through heterozygote advantage (Curtsinger and Heisler 1988; Fromhage et al. 2009), negative frequency dependence in host-parasite cycles (Hamilton and Zuk 1982) and antagonistic pleiotropy (Radwan et al. 2016; Li and Holman 2018), for example stemming from antagonistic selection in different environments.

In addition, spatially variable selection has been widely suggested to maintain V_a_ given limited gene flow between diverged populations (Hedrick 1986; Kisdi 2001; Byers 2005; Gray and McKinnon 2007; McDonald and Yeaman 2018). Specifically, given local adaptation, immigrants to any focal subpopulation will likely be maladapted and hence increase V_a_ in fitness (Lenormand 2002). Costly female preferences for local male displays could then be maintained by recurring immigration (Day 2000; Proulx 2001; Reinhold 2004). This mechanism could be commonplace if mating preferences and resulting sexual selection vary spatially (Payne and Krakauer 1997; Day 2000; Brooks 2002; Kingston et al. 2003; Chunco et al. 2007; Gray and McKinnon 2007; M’Gonigle et al. 2012; Wellenreuther et al. 2014; Ponkshe and Endler 2018; Dytham and Thom 2020). Indeed, several models have shown that evolution of divergent female preferences can cause spatially variable sexual selection, for example given spatially varying natural selection on male display (Lande 1982; Day 2000), varying male dispersal depending on mating success (Payne and Krakauer 1997) and spatial variation in carrying capacity combined with mate-search costs in females (M’Gonigle et al. 2012).

However, spatially varying natural selection or ecology may not be necessary for spatial processes to facilitate evolution of costly female preferences. Rather, spatially divergent preferences might evolve simply given initially neutral spatial population structure, defined here as spatial genetic variation arising from population subdivision with limited gene flow, without any ecological heterogeneity. Lande (1981) and Kirkpatrick (1982) both highlighted the potential importance of genetic drift in initiating preference-display coevolution, and later models showed that drift can generate divergence in mating preferences along the line of equilibrium between two completely isolated populations (Tazzyman and Iwasa 2010), even when preferences are costly (Uyeda et al. 2009). However, most models assumed infinite population size and constant genetic (co)variances (Mead and Arnold 2004; Kuijper et al. 2012), thus ignoring stochastic effects arising in finite populations. Therefore, the potential consequences of limited gene flow in facilitating persistence of V_a_ underlying display and associated preference evolution over long biological timeframes remain largely unexplored. Yet, within spatially structured populations, gene flow and genetic drift are both prominent processes which could shape G-matrices of sexual traits (Guillaume and Whitlock 2007; Dytham and Thom 2020; Reid and Arcese 2020), and thereby modulate costly preference and display coevolution.

Furthermore, although sexual selection is predicted to cause male display phenotypes to diverge from naturally selected optima (Lande 1981), the population dynamic consequences of resulting increased male mortality are seldom explicitly considered (Tanaka 1996; Kokko and Brooks 2003; Martínez-Ruiz and Knell 2017). Strong sexual selection might increase risk of evolutionary “suicide” (Kokko and Brooks 2003), limiting population persistence and hence inevitably eliminating costly display and preference genotypes. Such extinction risk might be greatest in small populations, where genetic drift is most prevalent. Overall temporal dynamics of preference-display coevolution might then differ markedly between spatially structured populations and large panmictic populations, as are typically considered.

To test our hypothesis that genetic drift, coupled with limited gene flow, can facilitate evolution and persistence of costly female preference and costly male display over long timeframes, we built a spatially and genetically explicit individual based model that allows genetic (co)variances of both traits and subpopulation extinction dynamics to arise from emerging spatial population structure. Specifically, we examine the consequences of dispersal probability and level of metapopulation subdivision for the temporal dynamics of preference and display coevolution and subpopulation extinction. We focus on temporal dynamics occurring over long biological timeframes, not specifically on evolutionary equilibria (stable states). Such transient persistence of preference-display co-evolution could still have considerable ecological and evolutionary consequences, even if V_a_ is ultimately completely depleted. We ask 1) whether dispersal among subpopulations can prolong the persistence of V_a_ in male display given uniform natural selection, and hence whether stochastic processes can be sufficient to fuel evolution and long-term persistence of costly female preference via indirect selection; 2) whether the degree of metapopulation subdivision interacts with dispersal to affect preference-display coevolution; 3) whether the degree and temporal trajectory of divergence in sexual traits depends on the level of spatial population structure; and 4) whether increasing spatial population structure elevates the risk of subpopulation extinction due to sexual selection. Consequently, we highlight the roles that initially neutral genetic variation and limited gene flow could play in shaping temporal evolutionary dynamics of sexual selection in spatially structured populations.

## Methods

We model a species inhabiting a square spatial grid where each cell contains a subpopulation connected by dispersal. We assume ecologically homogeneous space, where each cell has identical carrying capacity *K* and viability selection. At each non-overlapping generation, individuals undergo a lifecycle consisting of mating and reproduction, adult death, offspring viability selection and density-dependent survival, followed by dispersal. All model variables and parameters are summarised in Table S1.

### Genetic architecture

We model a diploid additive genetic system with two autosomal traits: female preference (*P*) and male display (*D*). Each individual carries L=10 diploid, physically unlinked loci underlying each trait (i.e., free recombination), with sex-limited phenotypic expression. Any genetic covariance between female and male traits arises from linkage disequilibrium generated by non-random mating. We assume a continuum-of-alleles model (Kimura 1965), with infinite potential alleles at each locus producing continuous distributions of effects. Initial allele values are independently sampled from a normal distribution with mean *θ_t_* (denoting the trait’s naturally selected optimum) and variance *σ^2^_α,0_* for both traits. Each individual’s genotypic value (*gP, gD*) is the sum of its 2L allele values. Phenotypes (*P* and *D*) correspond to the genotypes (i.e., no environmental variance), except we set *D*=0 if *gD*<0, meaning that male phenotypic display cannot be negative (e.g., envisaged as male crest or tail length), while female preference can take any real number.

### Mating and Reproduction

Starting each generation, each female chooses a mate from all N_males_ males present within her subpopulation according to her preference phenotype. Specifically, each male *j* has probability *p_j_* of mating with female *i*, which depends on the strength of female *i*’s preference (*P_i_*) and male *j*’s display (*D_j_*) relative to the displays of all other males in the subpopulation (Lande 1981; Bocedi and Reid 2015):

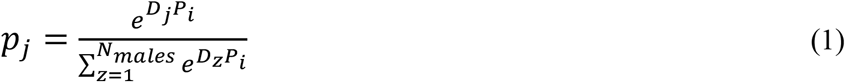

We model a psychophysical preference function where negative and positive values of *P* represent preference for smaller and larger than average displays respectively (Lande 1981). All females mate once, while males can mate multiple times or remain unmated. Each female then produces a number of offspring drawn from a Poisson distribution with mean *R*=4, assuming a 1:1 primary sex ratio. For each locus, each offspring inherits a randomly selected allele from each parent. Each allele has mutation probability of *μ*=5×10^-4^ per generation. Mutational effects are sampled from a normal distribution with mean *μ_m_* and variance *σ^2^_α_* (Table S1).

### Survival and Dispersal

Offspring survival results from consecutive viability selection and density-dependence. First, each offspring *i* experiences stabilising selection around a naturally selected optimum phenotype, with viability *v_i_* (Lande 1981; Bocedi and Reid 2015):

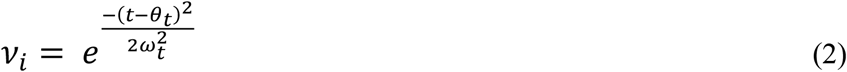

Here, *t* denotes individual *i*’s phenotype (i.e., *P* or *D*), *θ_t_* the trait’s naturally selected optimum (constant *θ_t_*=0 across subpopulations) and *ω_t_* the strength of stabilising selection (higher values give weaker selection). Any deviation from *θ_t_* due to sexual selection decreases the individual’s viability, imposing an absolute cost on preference or display. This may represent an energetic cost of male display (Basolo and Alcaraz 2003) or a “choosiness” cost of female preference (Jennions and Petrie 1997). After viability selection, each offspring survives with density-dependent probability (*ς*):

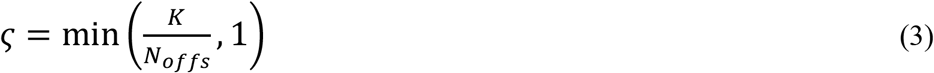

where N_offs_ is the subpopulation total number of offspring. The relative contributions to mortality of selection versus density-dependence thus depend on the distance of the subpopulation mean from the trait’s naturally selected optimum and the local subpopulation density after selection.

Each offspring that survives viability selection and density-dependent mortality may disperse to a different subpopulation with probability *d*. Distance and direction for each dispersal event are drawn from negative exponential (mean=1 cell) and uniform real (0-2π) distributions respectively, applied from a random starting point within the natal cell. If the destination falls outside the grid or within the current cell, distance and direction are resampled. This typically generates relatively short distance dispersal, with infrequent longer distance movements (Bocedi and Reid 2017).

### Simulation experiments

We investigated effects of spatial population structure on coevolution of female preference and male display in two ways. First, we examined effects of dispersal among subpopulations by varying dispersal probability (*d*=5×10^-5^,5×10^-4^,5×10^-3^,0.05) on a 7 by 7 cell grid (49 cells, each with *K*=154). Exploratory simulations showed that *d*>0.05 generated dynamics indistinguishable from a single panmictic population. Second, for each value of *d*, we varied the level of population subdivision by dividing a metapopulation totalling 7500 individuals into 9, 25, 49, 100 and 144 subpopulations of equal size (*K*=834, 300, 154, 75, 52 respectively).We assumed relatively weak stabilizing selection on female preference (ω^2^_P_=100), but strong stabilizing selection on male display (ω^2^_D_=4).

To evaluate effects of spatial population structure, we compared all scenarios to a single panmictic control population of 7500 individuals. This size was chosen to minimise effects of genetic drift while retaining reasonable computation time. Indirect selection favouring female preference evolution implies that mean *P* evolves to higher values than expected under mutation-(natural)selection-drift-migration balance. Accordingly, to distinguish effects of drift and indirect sexual selection, we ran simulations for each scenario of spatial population structure where all females mate randomly (i.e., no preference) but experience the same stabilising selection on a neutral trait acting as a control phenotype.

Each simulation was run for 250,000 generations, representing a long biological timeframe, and replicated 50 times. For each subpopulation, we extracted the mean trait genotypic values, genotypic variance, and the genetic correlation between preference and display at regular time intervals. We then calculated the grand mean across subpopulation genotypic means, variance and genetic correlations for each replicate and time point (hereafter subpopulation grand mean, mean subpopulation V_a_ and genetic correlation). We further report the standard deviation of subpopulation trait means per metapopulation as a measure of subpopulation divergence. At each recorded generation we also extracted the percentage of subpopulations per metapopulation that were extinct. For all results, we report the median and central 95% interval across replicates. Finally, to reveal underlying mechanisms driving the persistence of costly preference, we quantified the contributions of dispersal and segregation to V_a_ in display. Quantitative genetic models of the maintenance of costly preference invoke mutation or dispersal that is biased away from the locally preferred male genotypes (Pomiankowski et al. 1991; Day 2000). To test whether dispersal was biased in an analogous way, we quantified the degree of immigrant maladaptation resulting from spatially varying sexual selection (details in Supporting Information Fig. S5).

To test sensitivity to key parameters we ran additional simulations where the naturally selected optimum of male display *θ_D_*=10 (allowing females to prefer either bigger or smaller than optimal displays), with different strengths of stabilising natural selection on male display (ω^2^_D_=100) or female preference (ω^2^_P_=25, 400), lower mutation probability (*μ*=5×10^-6^), or more loci underlying each trait (*L*=100). We also ran simulations where all allelic values were initialised with zero for both traits to examine whether solely mutational input, alongside drift and spatial dynamics, could initiate preference-display coevolution.

## Results

### Low dispersal probabilities promote evolution of costly female preferences

Compared to a single panmictic population, simply restricting dispersal probability *d*, and hence generating neutral spatial population structure, readily led to ongoing evolution of costly female preference and male display over long biological timeframes (Fig. 1). The magnitudes of these effects depended on the value of *d*. Given low values (*d*<5×10^-2^), mean subpopulation female preference initially increased away from the naturally selected optimum of no preference (Fig. 1A). This increase in preference was mirrored by increasing male display (Fig. 1B). Exaggeration of both traits proceeded faster given *d*=5×10^-3^, reaching maximum mean values within the initial 10,000 generations, but occurred more slowly given *d*=5×10^-4^ (Fig. 1A-B). Eventually, after 250,000 generations, costly female preference only persisted at very low dispersal probability (*d*=5×10^-4^). Depletion of V_a_ in both traits was initially faster given lower *d*; however, the greatest V_a_ in display remained given dispersal probability *d*=5×10^-4^ (Fig. 1D).

**Figure 1.**
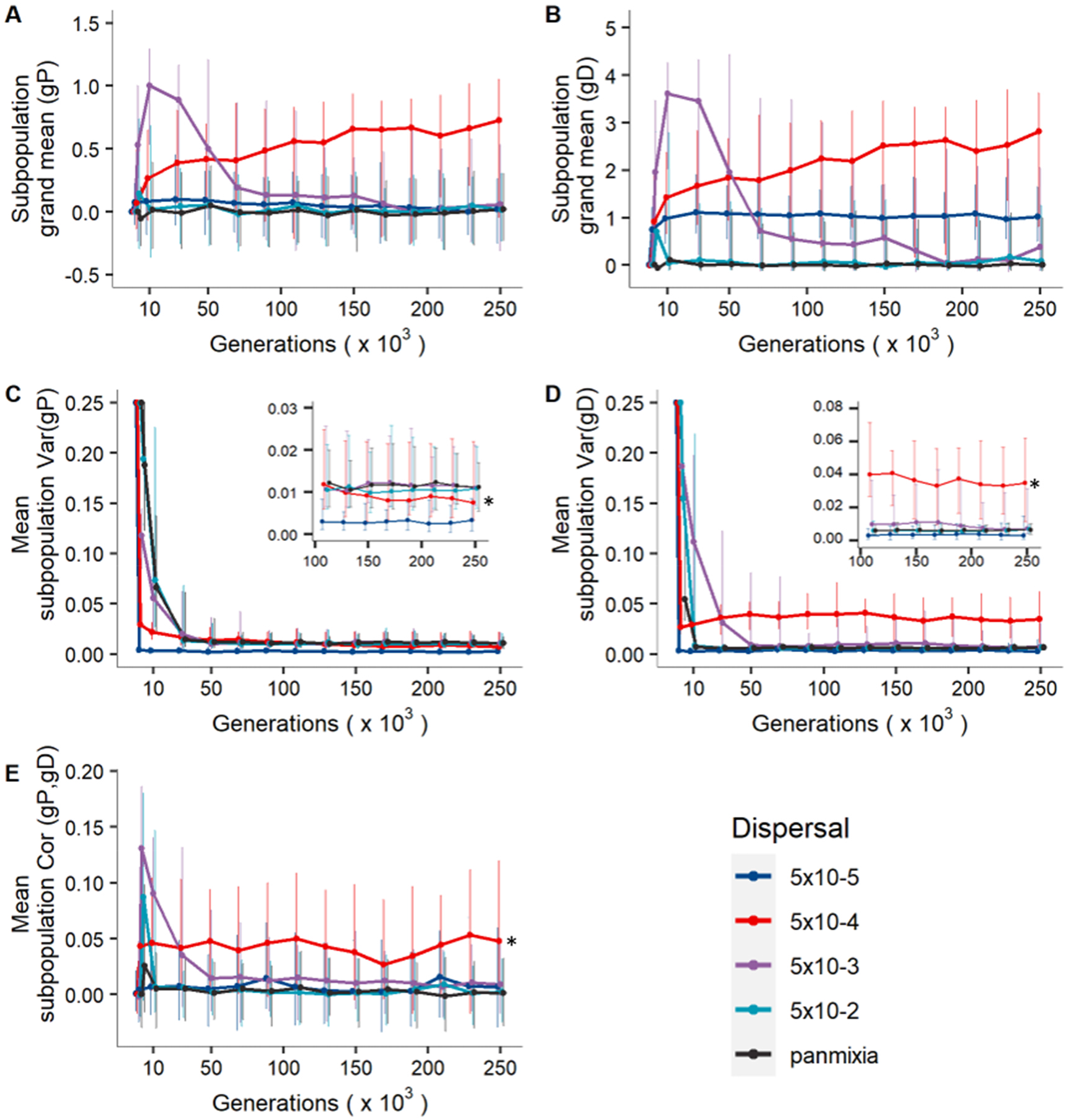
Coevolution of costly female preference and male display over 250,000 generations in metapopulations comprising 49 subpopulations given different dispersal probabilities (*d*). A) Subpopulation grand mean preference genotype *gP*; B) subpopulation grand mean display genotype *gD*; C-D) mean subpopulation additive genetic variance (V_a_) in *gP* and *gD*; F) mean subpopulation genetic correlation of preference and display Cor(*gP*, *gD*). In all panels, points and solid lines indicate the median across 50 replicate simulations, and bars indicate the central 95% interval across replicates. Data are shown initially at generation 0; 2,000; 10,000; and at intervals of 20,000 generations thereafter. Points are horizontally jittered to improve readability. Other parameters: ω^2^_P_=100 and ω^2^_D_=4. * indicates long-term transient dynamics.

A positive genetic correlation between preference and display initially arose given *d*>5×10^-4^ and peaked within the first 2,000 generations before gradually decreasing to zero (Fig. 1E). However, given *d*=5×10^-4^, a small positive genetic correlation persisted throughout the simulated period (Fig. 1E). In comparison, in control simulations with random mating, the distributions of genetic correlations were centred at zero, independent of dispersal probability (Fig. S1). Further, the distribution of subpopulation grand mean trait values was centred on the naturally selected optima. The difference between the main and control simulations (i.e., with and without sexual selection) suggests that indirect selection (in addition to drift) shaped preference evolution given preferential mating (Fig. S1).

Further analyses showed that dispersal between subpopulations was the largest source of V_a_ in male display (Fig.S4). In simulations where costly preference persisted, male immigrants were on average maladapted to the sexually selected environment at their destination, gene flow was therefore biased away from locally preferred male phenotypes (i.e., “migration bias”; Fig.S5).

Both short- and long-term persistence of female preference and male display, and hence the form of transient dynamics, depended on the strength of selection against preference (Fig. S6). Stronger selection (ω^2^_P_=25) prevented preference evolution irrespective of *d*. Conversely, weaker selection (ω^2^_P_=400) allowed transient increases in subpopulation grand mean preference even given *d*=0.05 (Fig. S6). Under weaker negative natural selection on male display (ω^2^_D_=100), display generally evolved to higher values compared to the main simulations (Fig. S8). Subpopulation grand mean preference and display showed pronounced fluctuations across generations given *d*<5×10^-4^, where preferences gradually changed from positive to negative, with corresponding responses in display (Fig. S8). Allowing evolution of either positive or negative male display by setting *θ_D_*=10 led to higher V_a_ in display at *d*=5×10^-4^, but preference and display evolution was otherwise equivalent to the main simulations (Fig. S10).

Results remained qualitatively similar given *L*=100 loci underlying preference and display (Fig. S12-S13). Lower mutation rate (*μ*=5×10^-6^) resulted in transient dynamics that were qualitatively similar to the main simulations, but complete depletion of V_a_ ultimately occurred within the simulated time period for all values of *d* (Fig. S15). Further, presence or absence of initial standing genetic variation in preference and display did not affect system state after 250,000 generations with *μ*=5×10^-4^ (Fig. S17). However, with lower mutation rates (*μ*<5×10^-4^), solely mutational input was not sufficient to initiate preference-display coevolution.

### Metapopulation subdivision and dispersal interact in determining preference-display coevolution

Increasing or decreasing the number of subpopulations (i.e., metapopulation subdivision) altered the evolution and persistence of female preference and male display depending on dispersal probability (Fig. 2A-B and Fig. S19-S22). The main effect was increasing long-term subpopulation grand mean preference and display with increasing metapopulation subdivision given *d*=5×10^-3^. In contrast, trait means decreased given *d*=5×10^-4^, and even turned negative given *d*=5×10^-5^ (Fig. 2A-B). With *d*=5×10^-5^, decreasing metapopulation subdivision steadily increased V_a_ in both preference and display (Fig. 2C-D). In contrast, at *d*=5×10^-4^ and 5×10^-3^, V_a_ was greatest given intermediate and high levels of metapopulation subdivision, respectively (Fig. 2C-D). Under random mating, subpopulation grand trait means remained at their naturally selected optima independently of metapopulation subdivision (Fig. S2). Further, greater V_a_ in display remained at *d*=5 × 10^-4^ and 5×10^-3^ given sexual selection than under random mating (Fig. S2).

**Figure 2.**
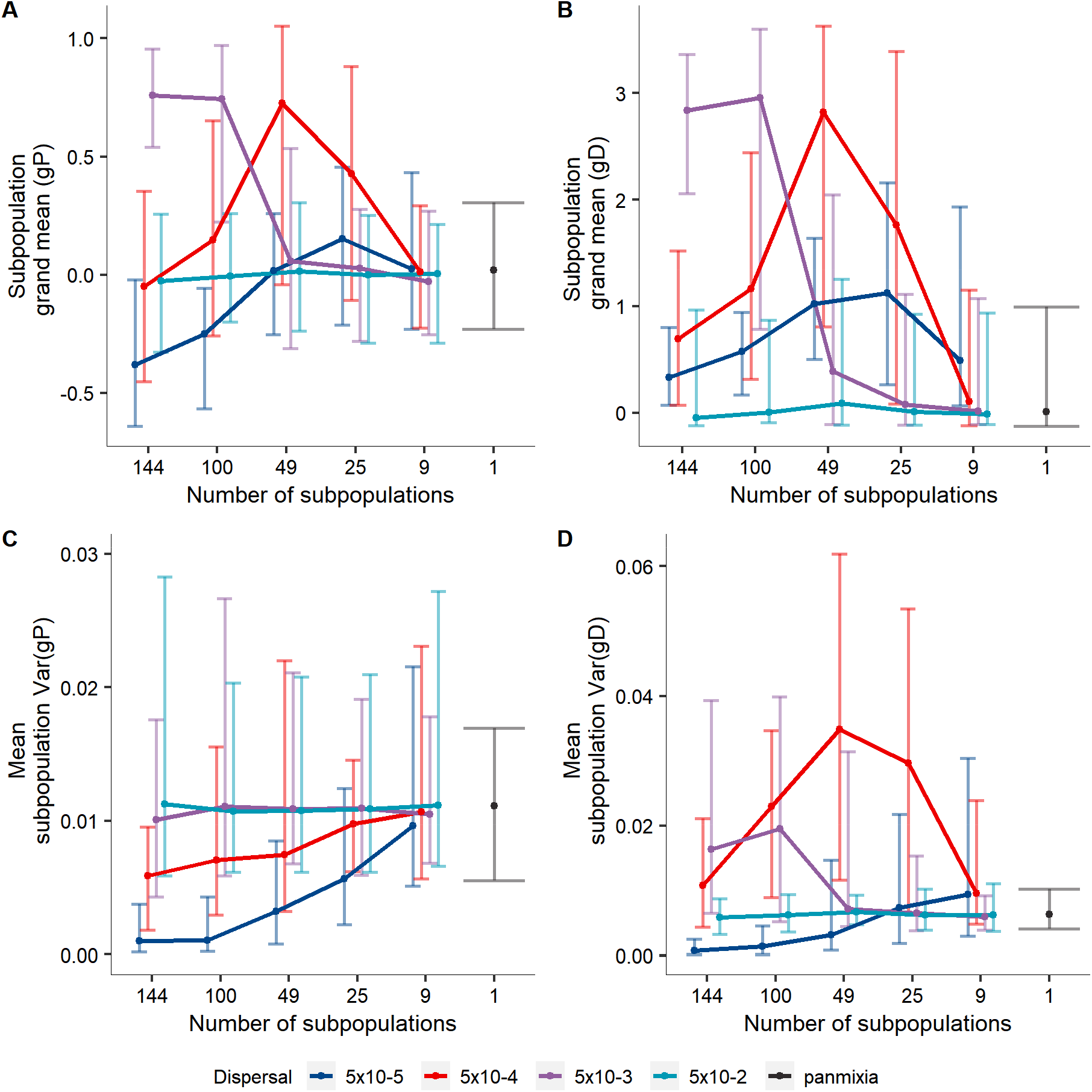
Effect of metapopulation subdivision (number of subpopulations) on the coevolution of costly female preference and male display, given different dispersal probabilities. A) Subpopulation grand mean preference genotype *gP*; B) Subpopulation grand mean display genotype *gD*; C-D) mean subpopulation additive genetic variance (V_a_) in *gP* and *gD*. In all panels, points and solid lines indicate the median across 50 replicate simulations at generation 250,000, and bars indicate the central 95% interval across replicates. Points are horizontally jittered to improve readability. Other parameters: ω^2^_P_=100 and ω^2^_D_=4.

### Effect of spatial population structure on subpopulation divergence

The levels of dispersal and metapopulation subdivision influenced the build-up and persistence of divergence in preference and display among subpopulations (Fig. 3). Under intermediate metapopulation subdivision (49 subpopulations), the lowest dispersal probability (*d*=5×10^-5^) resulted in high spatial differentiation in mean preference and display phenotypes across 250,000 generations, while *d* ≥5×10^-4^ led to short transient differentiation followed by subsequent homogenisation. Homogenisation occurred faster, and was complete across subpopulations, with higher *d* (Fig. 3A-B) and lower metapopulation subdivision (Fig. S23). However, with *d*=5×10^-4^ and high metapopulation subdivision, homogenisation proceeded very slowly, leaving substantial differentiation even after 250,000 generations (Fig. 3C-D, Fig. S23). Population differentiation in preference generally increased with increasing subdivision, while differentiation in display was highest at intermediate subdivision (Fig. 3C-D). In contrast, simulations with random mating (i.e., no sexual selection) showed less population differentiation, especially in male display (Fig. S3).

**Figure 3.**
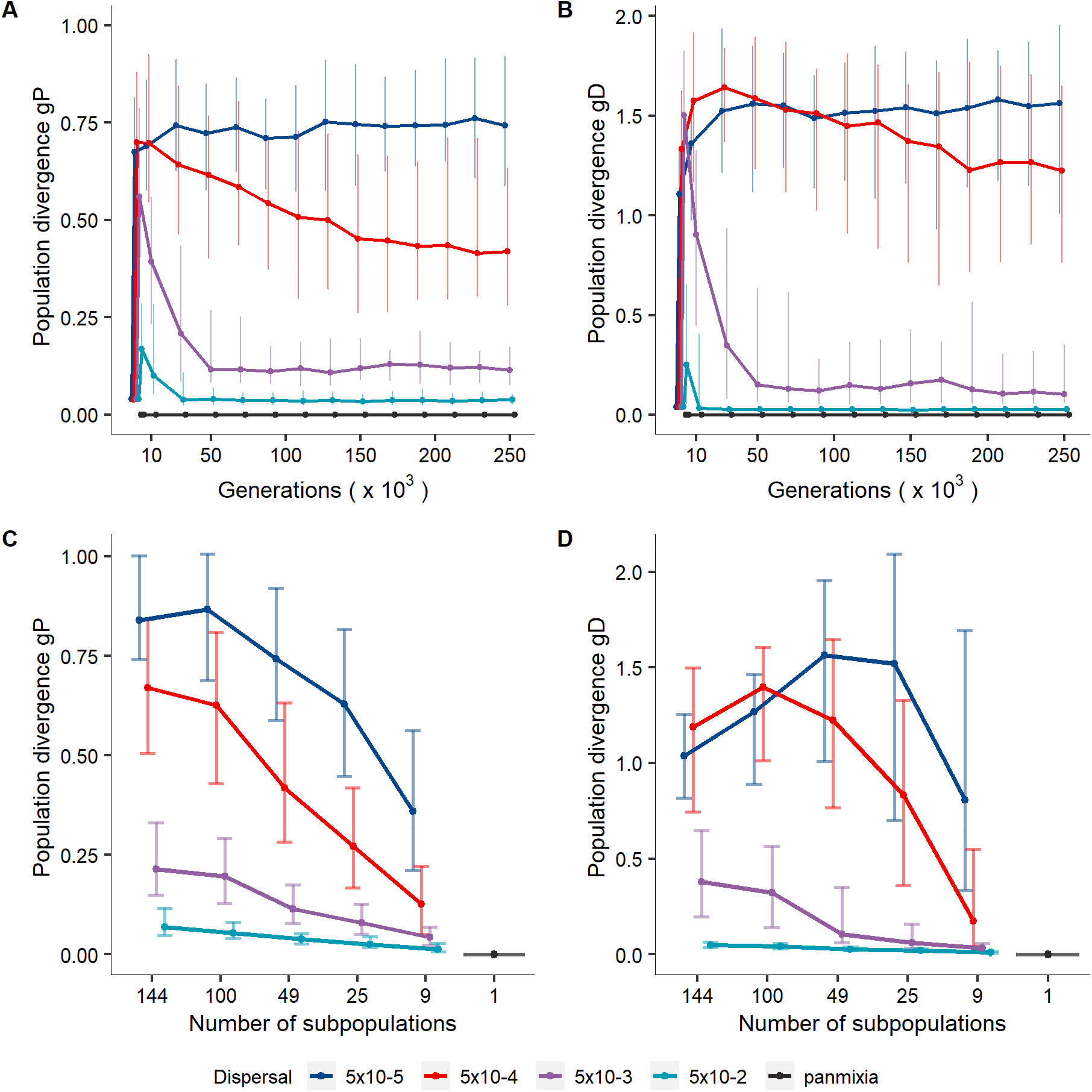
Population divergence in female preference genotype *gP* and male display genotype *gD* (A-B) over 250,000 generations and (C-D) at different levels of metapopulation subdivision (number of subpopulations) as a function of dispersal probability. Subpopulation divergence is measured as standard deviation of the subpopulation trait means across a metapopulation. A-B consider metapopulations composed of 49 subpopulations and data are shown initially at generation 0; 2,000; 10,000; and at intervals of 20,000 generations thereafter. C-D shows results at generation 250,000. In all panels, points and solid lines indicate the median across 50 replicate simulations, and bars indicate the central 95% interval across replicates. Points are horizontally jittered to improve readability. Other parameters: ω^2^_P_=100 and ω^2^_D_=4.

Weaker direct selection against male display resulted in higher and fluctuating population differentiation in display given *d* ≤5×10^-4^, but faster homogenisation given *d* >5×10^-4^ (Fig. S9). Setting optimal display phenotype *θ_D_*=10 resulted in greater population differentiation in both preference and display, because preference-display coevolution proceeded towards both smaller and larger than optimal display, depending on the subpopulation (Fig. S11). The degree of population differentiation remained similar with L=100 loci underlying the traits (Fig. S14).

### Increasing spatial population structure leads to subpopulation extinction via sexual selection

Evolution of mating preference (i.e., sexual selection) led to subpopulation extinction events to degrees that depended on spatial population structure (Fig. 4). With sexual selection, some subpopulation extinction occurred given *d*<5×10^-2^ and metapopulation subdivision >25 subpopulations (Fig. 4). Unsurprisingly, increasing subdivision (i.e., decreasing subpopulation size) generally increased the proportion of subpopulations that went extinct (Fig. 4), occasionally resulting in extinction of entire metapopulations (Fig. S24). In contrast, no extinctions occurred with random mating, indicating that extinctions were driven by sexual selection.

**Figure 4.**
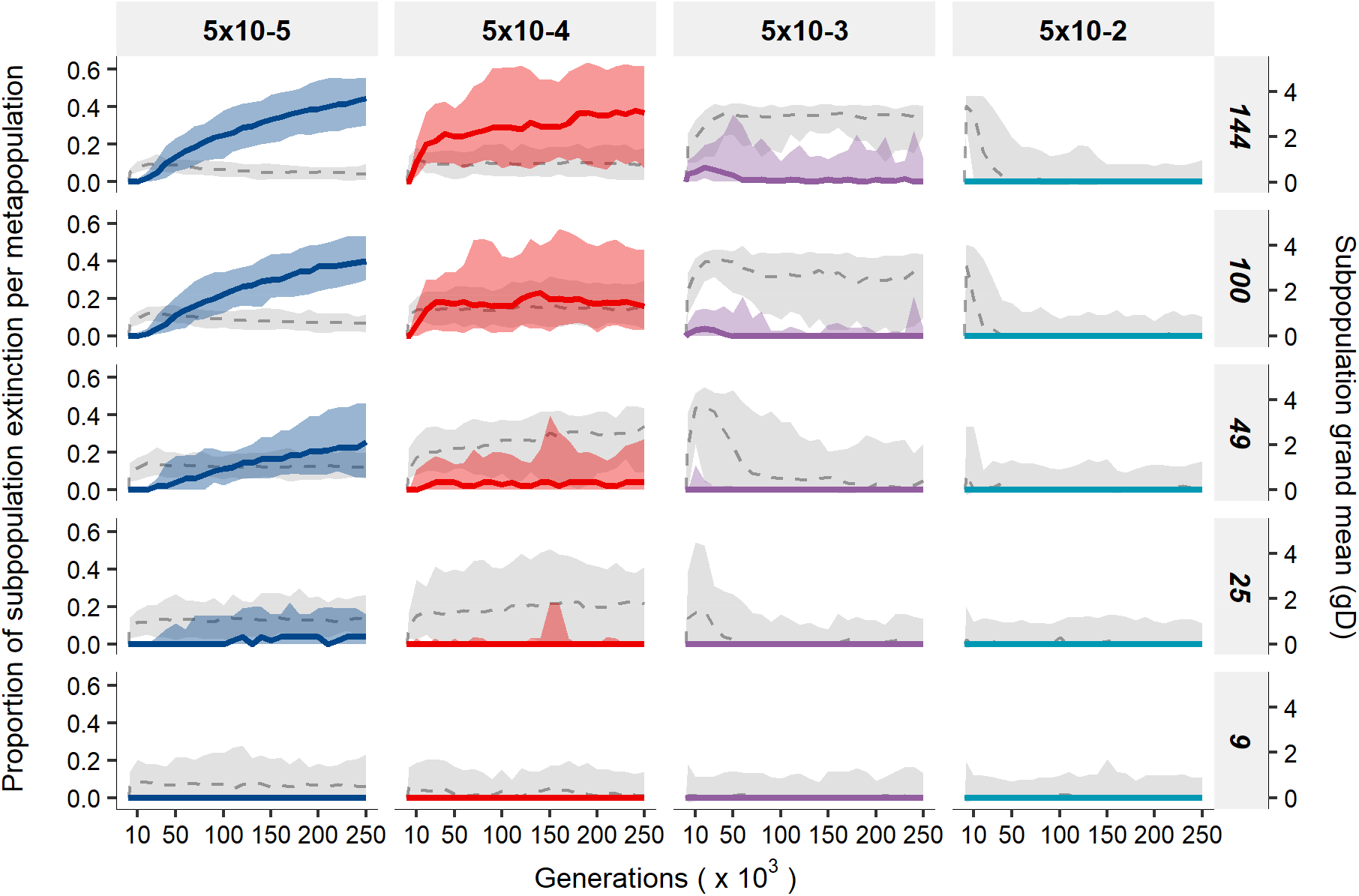
Proportion of subpopulations that went extinct (left y-axis) and male display evolution (right y-axis) over 250,000 generations, across different dispersal probabilities and levels of metapopulation subdivision. Colours and solid lines indicate population extinction, while dashed lines and grey shading refer to subpopulation grand mean display genotype *gD*.Lines indicate the median across 50 replicate simulations, and shading indicate the central 95% interval across replicates. Grey panels indicate different simulation scenarios of spatial population structure (x-axis: dispersal probabilities; y-axis: metapopulation subdivision).

## Discussion

Previous stochastic models of coevolving female preference and male display provide important insights into interactions between sexual selection and genetic drift (Uyeda et al. 2009; Tazzyman and Iwasa 2010). These studies demonstrate diversification of mating preferences under drift and examine the consequences for evolution of preferred traits in the absence of gene flow. Here, we extend these concepts to examine the combined effects of genetic drift and dispersal in shaping preference-display coevolution in a metapopulation context within an ecologically homogeneous environment. Specifically, given low dispersal probability, drift-induced female preference evolution created spatially variable sexual selection. Subsequent immigration of male genotypes that were maladapted to the local sexually selected environment then led to a “migration bias” (Day 2000) that created V_a_ in male display. Consequently, females exhibiting mating preference continued to produce more attractive (fitter) sons, thereby fuelling persistence of costly female preference across very long biological timeframes. Hence, initially neutral spatial population structure prolonged the persistence of V_a_ in male display compared to a panmictic population of identical overall size. However, these long transient dynamics of display and preference occurred only under a narrow range of low dispersal probabilities. Drift-induced evolution of costly female preference and costly male display was notably also associated with increased subpopulation extinction.

Our work further emphasises the need to consider sexual selection in a spatially explicit ecological context to understand the evolution and persistence of sexual traits (Payne and Krakauer 1997; Day 2000; M’Gonigle et al. 2012). A recent study examined effects of population fragmentation on the diversity of individual identity signals under sexual selection, and found that global signal diversity increased by 10-15% in fragmented versus unfragmented habitat (Dytham and Thom 2020). Our results similarly suggest that fragmentation (here represented by strong spatial structure) elevates genetic variation in male sexual signals and extensively prolongs the persistence of costly female preference. This raises the interesting empirical question of whether ongoing anthropogenic habitat fragmentation might ultimately elevate evolution and differentiation in species’ sexually selected traits. Most empirical evidence of sexual trait differentiation as a consequence of habitat fragmentation relates to alteration of biotic and abiotic environments, such as changes in predation pressures or water turbidity (Giery et al. 2015; Giery and Layman 2015; Zastavniouk et al. 2017). Yet, population differentiation in mating preferences that is not clearly associated with such environmental changes could indicate that the genetic consequences of habitat fragmentation have fuelled preference evolution. Interestingly, increasing habitat fragmentation may increase costs of dispersal (Bonte et al. 2012; Cote et al. 2017), selecting against it and thereby facilitating preference-display coevolutionary dynamics under low dispersal.

In our model, greatest V_a_ in male display persisted over long timeframes given low dispersal probabilities (*d*=5×10^-4^ and 5×10^-3^). We showed that dispersal between subpopulations is the main cause, with very low contribution from mutation and segregation (Fig.S4). This indicates that prolonged persistence of V_a_ is tightly linked to the level of subpopulation differentiation in display. While high metapopulation subdivision promoted differentiation, such systems require more dispersal to counteract rapid depletion of V_a_ due to smaller subpopulations size. Ultimately, because our model allowed the source of spatial variation in sexual selection (i.e., female preference) to evolve, homogenisation of female preference with increasing dispersal generally caused depletion of V_a_ in display.

Our results generally concur with recent findings that Fisherian sexual selection alone cannot maintain population divergence given substantial gene flow (Servedio and Bürger 2014, 2015). Servedio and Bürger (2014) examined divergence in a fixed relative preference and a male display trait in a two-locus haploid model, and found that increasing sexual selection inhibited differentiation between populations connected by gene flow due to limited divergence in preference. When female preference could evolve, weaker preference alleles successfully invaded until preference disappeared. However, at low dispersal, our analysis of quantitative variation underlying preference and display showed that gene flow played a crucial role in generating a positive genetic correlation across initial generations, and divergence in sexual traits often persisted for substantial periods of time in structured populations. Similarly, M’Gonigle et al. (2012) showed how variation in local carrying capacity combined with mate-search dependent costs to females can facilitate prolonged persistence of ecologically equivalent mating types. In our model, similar persistence of variation in preference and display emerged with neither variation in local carrying capacity nor mate-search costs.

Drift-induced evolution of female preference has been shown to increase differentiation between populations (Uyeda et al. 2009; Tazzyman and Iwasa 2010) and is expected to help initiate Fisherian preference-display coevolution (Lande 1981). However, genetic drift is higher in small populations, which also have higher extinction risk. Our simulations highlight this balance for small and fragmented populations: genetic drift and little dispersal promote evolution of female preference for costly male display, but such populations are also prone to extinction, likely because of increased mortality due to expression of costly male traits (Kokko and Brooks 2003; Martínez-Ruiz and Knell 2017). Further, under some scenarios, specifically with weak stabilising natural selection on preference and/or display, sub-population extinctions can lead to rapid shifts in the metapopulation trait means that somewhat resemble previously described cycles of preference-display coevolution (Iwasa and Pomiankowski 1995; Kuijper et al. 2012). In our case, trait changes are apparently driven by the combination of weak stabilising selection leading to runaway coevolution of the sexual traits, and associated extinction-recolonisation dynamics (cf. Fig. S8 and Fig. S25). Interestingly, some degree of population extinction may contribute positively to the persistence of costly preference since almost extinct populations will generate fewer migrants, contributing to biased dispersal towards individuals with lower display values than the metapopulation average (Fig. S26). This raises interesting questions regarding potential interactions between metapopulation dynamics and maintenance of V_a_ for sexual traits, which remain to be explored.

The observed local extinction dynamics presumably depend on assumptions regarding the strength and form of natural selection on display. We modelled strong enough natural selection to outweigh genetic drift even in highly structured populations, thereby highlighting evolution of costly display in response to sexual selection. We assumed natural selection to be relative to a global phenotypic optimum, independently from local male display genotypes and density (Reznick 2016; De Lisle and Svensson 2017). Alternative modes of selection would likely alter outcomes. For example, frequency dependent natural selection may reduce the costs of display locally, reducing male mortality and allowing subpopulations to diverge further compared to a global phenotypic optimum, with potentially positive effects on the persistence of V_a_. Conversely, higher population differentiation might increase the costs of display disproportionally for immigrant males if selection acts relative to the local mean phenotype, potentially limiting the persistence of V_a_ by reducing gene flow between subpopulations (Gosden et al. 2015). While our assumption about the form and strength of natural selection may particularly promote male mortality, this effect is likely counteracted by our assumption that all local females could reproduce with a single surviving male. Allee affects arising from limited mate availability may otherwise be expected to further increase population extinction risk (Shaw and Kokko 2014). Future studies could investigate different forms of natural selection on display and link costs of female preferences, in terms of missed reproduction opportunities, to local male density. Similarly, alternative assumptions regarding individual dispersal provide avenues for future investigations. Dispersal decisions and costs are complex, and often depend on individual phenotype and/or local competition (Bowler and Benton 2005). Introducing such complexity could cause further migration bias when males with larger displays pay proportionally higher costs of dispersing.

Overall, our model demonstrates that purely spatial population structure, arising from population subdivision and restricted ranges of limited dispersal, can fuel costly preference-display evolution and allow V_a_ in display to persist over long biological timescales without invoking spatially variable natural selection on display or unrealistically high mutation rates. Yet, we show that populations with extreme preference-display genotypes might experience evolutionary suicide, driving long-term extinction-recolonisation dynamics. These analyses should inspire future investigations of preference-display coevolutionary dynamics across more complex spatial and fitness landscapes.

