## Supporting Information for "Strong spatial population structure shapes the temporal coevolutionary dynamics of costly female preference and male display"

### **Table S1.** Summary of variables and parameter values used in models of coevolution of preference (*P*) and display (*D)*

|  | **Variables** | **Description** | **Parameter values** |
| --- | --- | --- | --- |
|  | *K* | subpopulation carrying capacity | 834, 300, 154, 75, 52 |
|  |  | number of subpopulations | 9, 25, 49, 100, 144 |
|  | *d* | dispersal probability | 5×10^-5^, 5×10^-4^, 5×10^-3^, 0.05 |
|  | *R* | mean fecundity | 4 |
|  | *L* | number of diploid loci for each trait | 10; 100 (sensitivity analysis) |
| **Mutation** | *µ* | mutation probability (per allele per generation) | 5×10^-4^; 5×10^-5^, 5×10^-6^ (sensitivity analysis) |
|  | *μ_m_* | mean mutational effect | 0.0 |
|  | *σ^2^_α_* | variance in mutational effects for both traits | = (0.25 /2*L*)*0.1 |
| **Traits** | *gP* | female preference for male display (genotypic value) |  |
|  | *gD* | male display (genotypic value) |  |
|  | *P* | female preference for male display (phenotypic value) | =$g$P |
|  | *D* | male display (phenotypic value) | =$\left\{ \begin{matrix} 0 & \mathrm{for} & gD<0 \\ D & = & gD \geq0 \end{matrix} \right.$  =$gD$ (sensitivity analysis) |
| **Trait Initialisation** | *µ_P,0_* | initial genotypic mean for female preference | 0.0 |
|  | *µ_D,0_* | initial genotypic mean for male display | 0.0; 10.0 (sensitivity analysis) |
|  | *σ^2^_G,0_* | initial genotypic variance for both traits | 0.25; 0 (sensitivity analysis) |
|  | *µ_aP,0_* | initial allelic mean for female preference | = *µ_P,0_* / 2*L* |
|  | *µ_aD,0_* | initial allelic mean for male display | = *µ_D,0_* / 2*L* |
|  | *σ^2^_a,0_* | initial allelic variance for the two traits | = *σ^2^_G,0_* / 2*L* |
| **Costs** | *θ_P_* | naturally selected optimum for preference | 0.0 |
|  | *θ_D_* | naturally selected optimum for display | 0.0; 10.0 (sensitivity analysis) |
|  | *ω^2^_P_* | strength of stabilising natural selection for preference | 100; 25, 400 (sensitivity analysis) |
|  | *ω^2^_D_* | strength of stabilising natural selection for display | 4; 100 (sensitivity analysis) |

### Simulations under random mating

To distinguish between the effect of drift and the effect of indirect sexual selection on female preference evolution, we ran simulations in which all females mate randomly but experience the same level of stabilising selection on a neutral trait (*FC*), acting as a control phenotype under mutation-(natural)selection-drift-migration balance. The distribution of subpopulation grand mean trait values across replicate simulations was centred on the naturally selected optima (Fig. S1A-B and Fig. S2A-B). While the maintenance of the genetic variation in the female control trait was the same as in our main analysis (Fig. S1C and Fig. S2C), genetic variation in male display depleted earlier under random mating (Fig. S1D) and was generally higher under preferential mating for dispersal probability of 5×10^-4^ and 5×10^-3^ (cf. Fig 2D with Fig. S2D). The distributions of genetic correlations between female control trait and male display were centred at zero, and independent of dispersal probability (Fig. S1E). The difference between the control simulations and our main results suggests the action of indirect selection (in addition to drift) in determining evolution of subpopulation preferences under preferential mating. Further, simulations under random mating showed slightly lower levels of population divergence in the female control trait and differentiation in male display was minimal (Fig. S3).

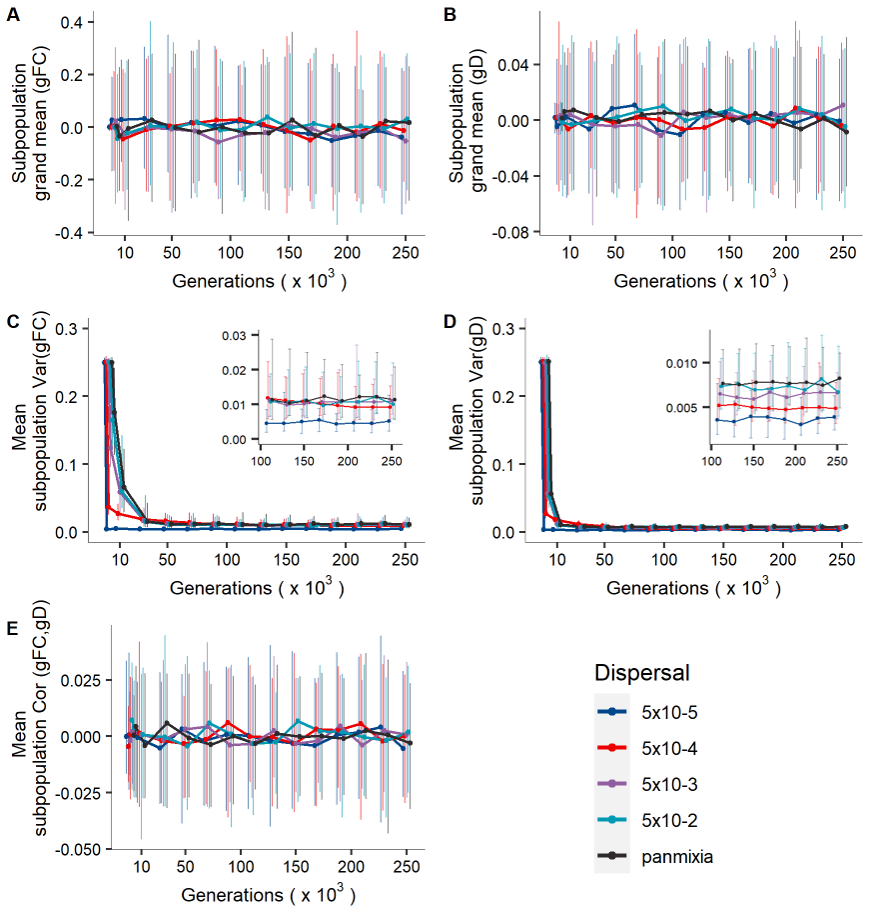

**Figure S1.** Coevolution of female control trait (FC, random mating) and male display in metapopulations composed of 49 subpopulations under varying levels of dispersal probability over 250,000 generations. Panels, axes and lines as in Figure 1.

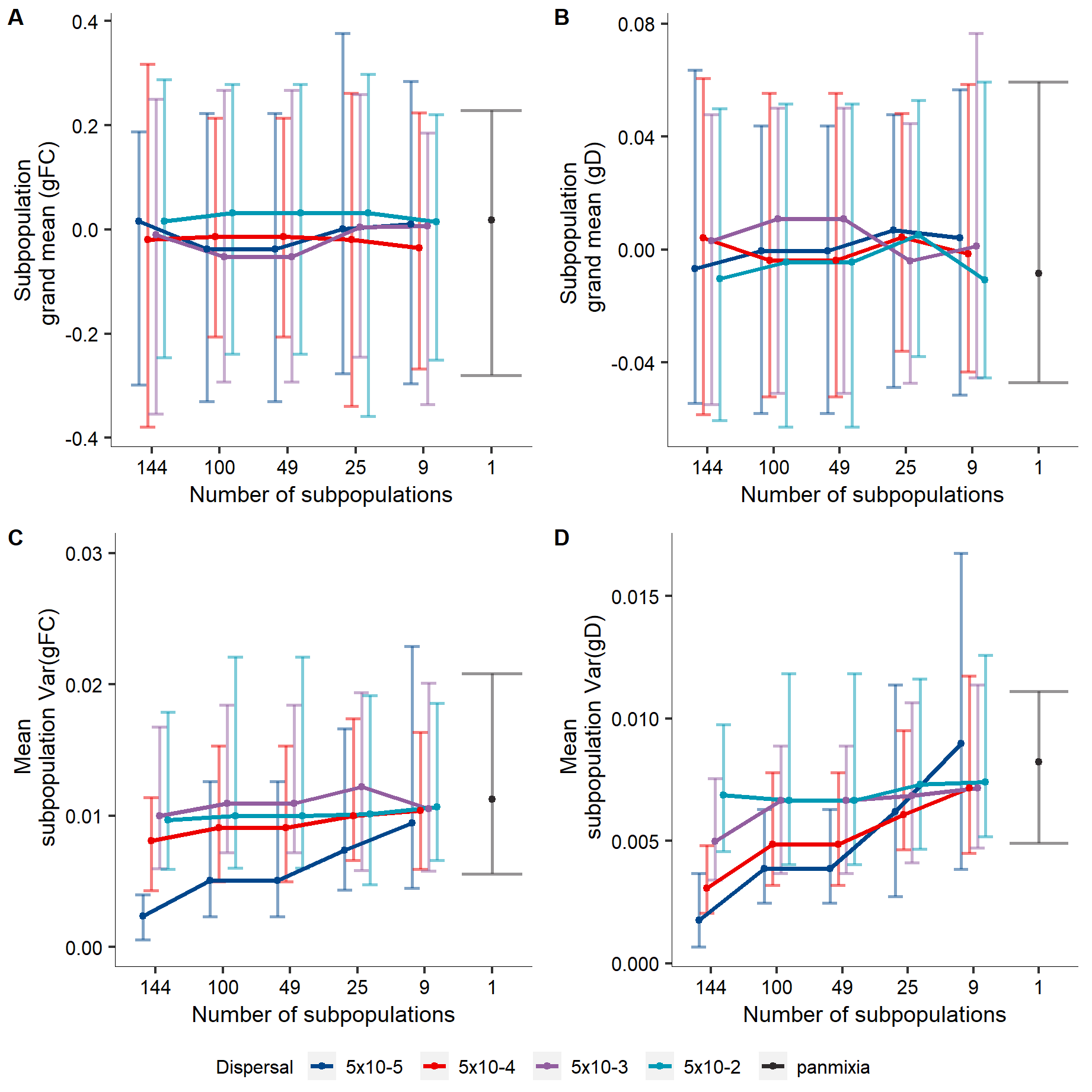

**Figure S2.** Effect of metapopulation subdivision (number of subpopulations) on the coevolution of a female control trait and male display, given different dispersal probabilities. Panels, axes and lines as in Figure 2.

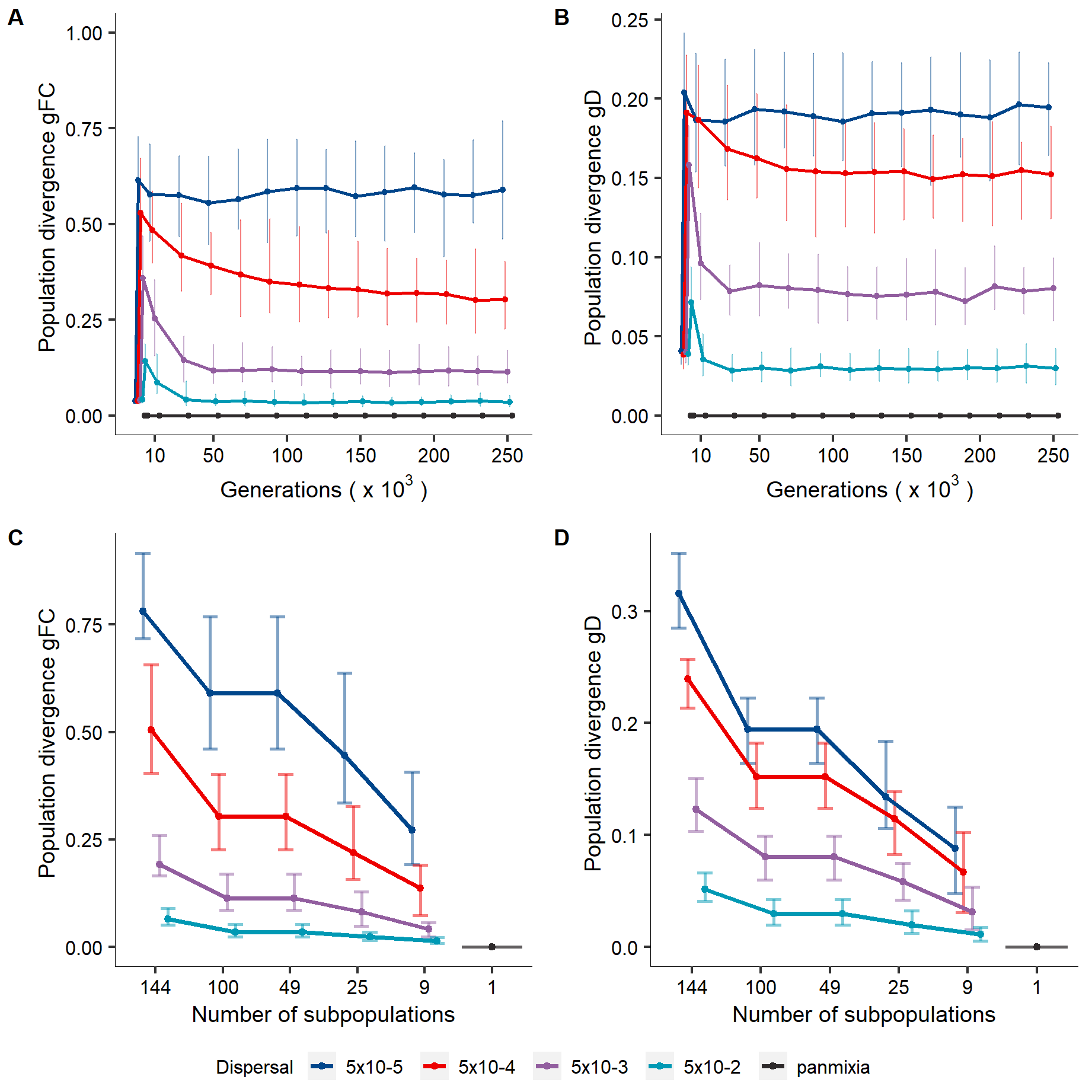

**Figure S3.** Population divergence in female control trait genotype *gFC* and male display genotype *gD* (A-B) over 250,000 generations and (C-D) with different levels of metapopulation subdivision (number of subpopulations) as a function of dispersal probability at generation 250,000. Panels, axes and lines as in Figure 3.

### Sources of variance in display

Persistence of additive genetic variance ($V_{a}$) in display given directional selection must result from a combination of mutation, dispersal and segregating variance. To reveal the mechanism underlying long term persistence of$V_{a}$ in display in our model, we quantified the relative contributions of dispersal (i.e., dispersal variance$V_{a_{D}}$) and segregation (i.e., segregating variance $V_{a_{S}}$) to $V_{a}$ in display. We additionally compare this to theoretical expectations about the degree of $V_{a}$ contributed by mutation (i.e., mutational variance$V_{a_{M}}$), which is expected to be $V_{a_{M}}= 2L\mu\sigma_{\alpha}^{2}$ (Walsh and Lynch 2019).

We define $V_{a_{D}}$ as the per generation contribution to $V_{a}$ arising from dispersal of individuals between subpopulations. We calculated $V_{a_{D}}$in display as $V_{a_{D}}=$ ${V_{a}}_{I+R}- {V_{a}}_{R}$, where ${V_{a}}_{I+R}$ is $V_{a}$ in *gD* calculated from residents and newly immigrated individuals ($V_{a}$ in *gD* after dispersal), and ${V_{a}}_{R}$ is the $V_{a}$ in *gD* calculated from only residents ($V_{a}$ in *gD* before dispersal excluding emigrants).

Further, we define$V_{a_{S}}$ as per generation contribution to $V_{a}$ arising from recombination in the offspring generation. As such, we could in principle calculate $V_{a_{S}}$ as the difference between the $V_{a}$ in *gD* after and before reproduction. However, preferential mating and genetic drift reduce the amount of $V_{a}$in *gD*, while recombination and mutation increase the amount of $V_{a}$ in the offspring generation. Accordingly, we computed a measure of $V_{a_{S}}$that excludes both the effect of mutation and preferential mating by randomly mating females with males, and subsequently increase the number of hypothetical offspring per female to 40 to reduce the magnitude of genetic drift. We then calculate $V_{a_{S}}$ in display as $V_{a_{S}} ={V_{a}}_{hypothetical offspring}- {V_{a}}_{adults}$ where ${V_{a}}_{adults}$ is $V_{a}$in *gD* calculated from adults before reproduction occurs, and ${V_{a}}_{hypothetical offspring}$ is $V_{a}$ in *gD* calculated from their hypothetical offspring. We then calculated the grand mean across subpopulation dispersal variance and subpopulation segregating variance for each replicate and time point.

In Figure S4 we show results for a metapopulation composed of 49 subpopulations, the same parameters as in our main simulations (Figure 1). Here, we show that dispersal variance $V_{a_{D}}$ provides the most important per-generation contribution to$V_{a}$ in display. For *d* = 5×10^-4^, mean subpopulation dispersal variance is ≈ 0.025, contributing on average 65% of the variance observed in display (compare Fig. S4A with Fig. 1D). In comparison, our approximation of segregating variance $V_{a_{S}}$ shows that its mean contribution is negligible (yet highly variable) and is regularly even slightly negative for *d* = 5×10^-4^ on average on the order of -5×10^-4^ (Fig. S4B). Negative $V_{a_{S}}$ indicates that some of the dispersal variance $V_{a_{D}}$arising in adults may be regularly lost in the offspring generation due to genetic drift arising from stochasticity in mate sampling. Additionally, the contribution of mutational variance $V_{a_{M}}$is expected to be also very small, on the order of 1.25×10^-5^.

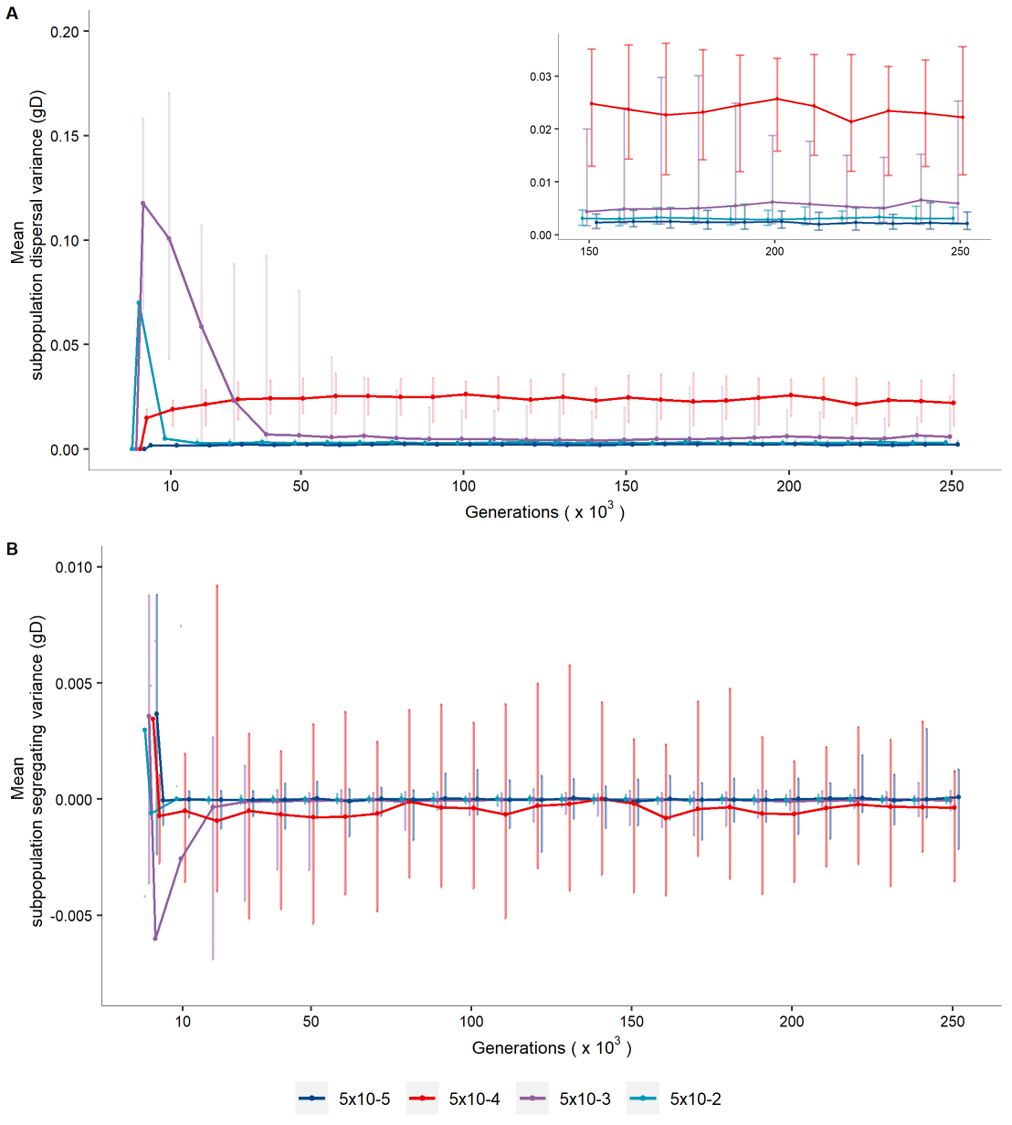

**Figure S4**. Contribution of dispersal variance (A) and segregating variance (B) to the total additive genetic variance in male display. Results are shown for metapopulations composed of 49 subpopulations, for different dispersal probabilities (colours). Axes and lines as in Figure 1.

.

### Immigrant maladaptation to the sexually selected environment

Maintaining a costly preference in quantitative genetic models requires the source of genetic variation to be biased towards the opposite direction of the locally preferred male genotypes (Pomiankowski et al. 1991; Day 2000). For dispersal variance to be biased in such a way, immigrants must on average be less attractive to the average female in the destination subpopulation, i.e., immigrant males must be maladapted to the sexually selected environment in their destination subpopulation. To show that dispersal variance is indeed biased in our model, we quantified the average maladaptation of male immigrants to the sexually selected environment in their destination subpopulation. We then calculated two measures of maladaptation: the immigrants maladaptation relative to the average male in the population, ${p_{I}}/{p_{A}}$ (where $p_{A}$ is the mating probability of the average male; Fig. S5A), and the immigrants maladaptation relative to the best (most attractive) male in the population, ${p_{I}}/{p_{B}}$ (where $p_{B}$ is the mating probability of the most attractive male; Fig. S5B). These two measures were calculated for each immigrant arriving in a subpopulation at a given generation, and then averaged across immigrants in the same subpopulation. We then averaged the subpopulation means at each generation to calculate subpopulation grand means across the entire metapopulation, obtaining the average mating probability of immigrants relative to the subpopulation average male, $\bar{p}_{IA}$, and the average mating probability of immigrants relative to the subpopulation most attractive male $\bar{p}_{IB}$. Because we model a directional open-ended preference, $\frac{p_{I}}{p_{B}}$ provides a better picture of actual maladaptation to the directional selection imposed. However, $\frac{p_{I}}{p_{A}}$ provides some intuition about male maladaptation with regards to the rest of the population.

In Figure S5 we show results for a metapopulation composed of 49 subpopulations and use the same parameters as in our main simulations (Figure 1). We show that the same dispersal scenarios that exhibit prolonged persistence of costly preference, also exhibit maladaptation of male immigrants to the sexually selected environment in their destination subpopulation. Such maladaptation of male immigrants therefore indicates a “migration bias” which prolongs the persistence of costly preference in our model.

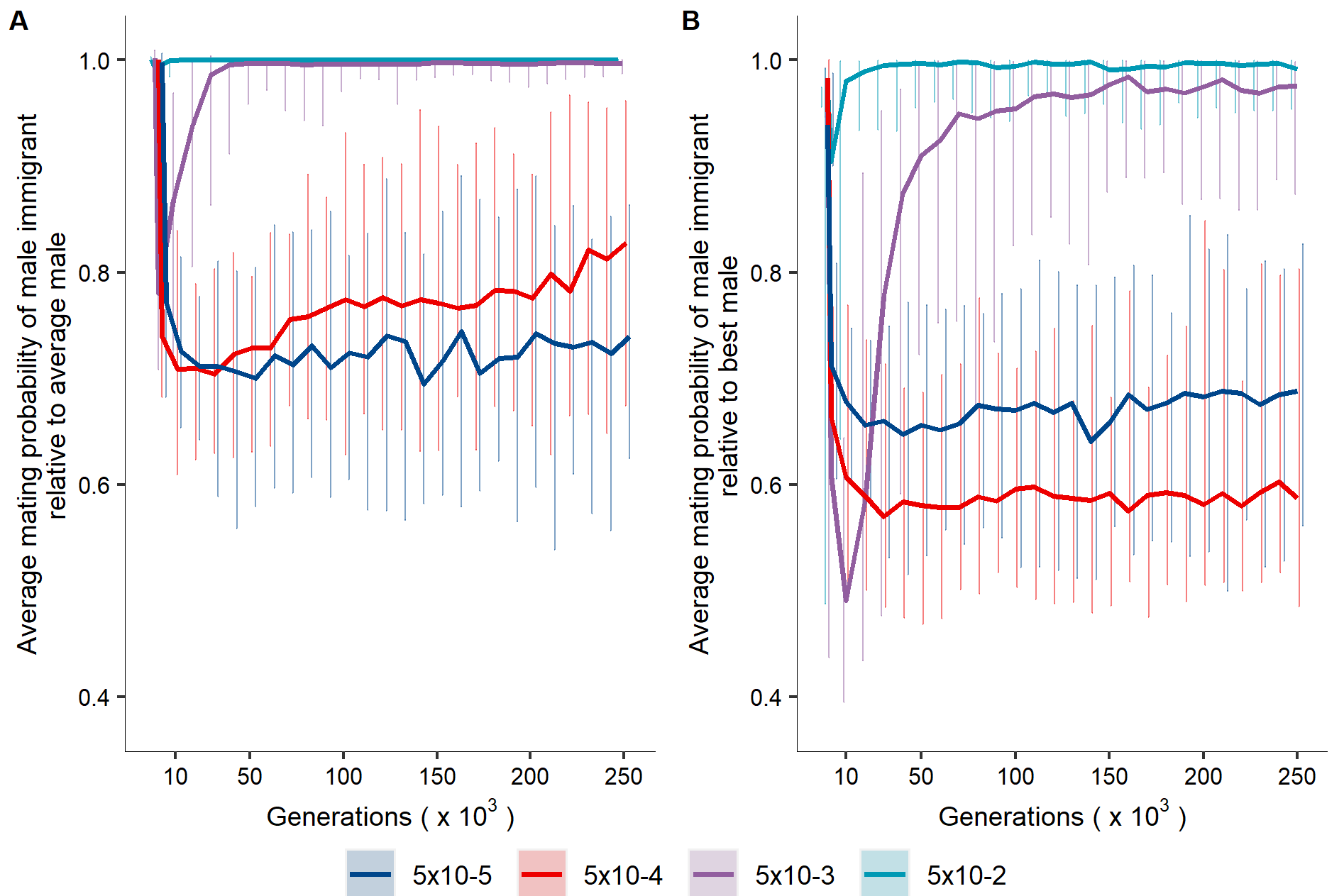

**Figure S5.** Two measures of immigrant maladaptation to the sexually selected environment in metapopulation composed of 49 subpopulations. A) Average mating probability of male immigrants relative to the average male, $\bar{p}_{IA}$; B) average mating probability of male immigrant relative to the best (most attractive) male, $\bar{p}_{IB}$. Axes and lines as in Figure 1.

### Effect of the strength of natural selection on female preference

We investigated the consequences of changing the strength of stabilising natural selection on female preference (ω^2^_P_ = 400, 25; i.e., weaker and stronger selection respectively). Increasing direct selection on female preference led the subpopulation grand mean preference to remain at its naturally selected optimum irrespective of dispersal (Fig. S6A, ω^2^_P_ = 25). Weaker selection on preference led to a prominent increase in the subpopulation grand mean preference under low-intermediate dispersal (Fig. S6A, ω^2^_P_ = 400, *d* = 5×10^-3^ and 5×10^-4^ ). Under *d* = 5×10^-5^, a negative preference evolved, whereby extinction of populations exhibiting particularly strong preferences may have contributed to shift the subpopulation mean preference towards negative values. For most dispersal probabilities, stronger and weaker selection on female preference led to similar levels of genetic variation in sexual traits as in our main analysis (ω^2^_P_ = 100), with the exception of lower V_a_ in display at *d* = 5×10^-4^ . While prominent divergence in sexual traits between subpopulations was observed under weak selection on female preference, strong selection reduced divergence (Fig. S7A-B). Nevertheless, low dispersal (*d* = 5×10^-4^ and 5×10^-5^) led to increasing population divergence in preferences under strong selection due to drift (Fig. S7A, ω^2^_P_ = 25).

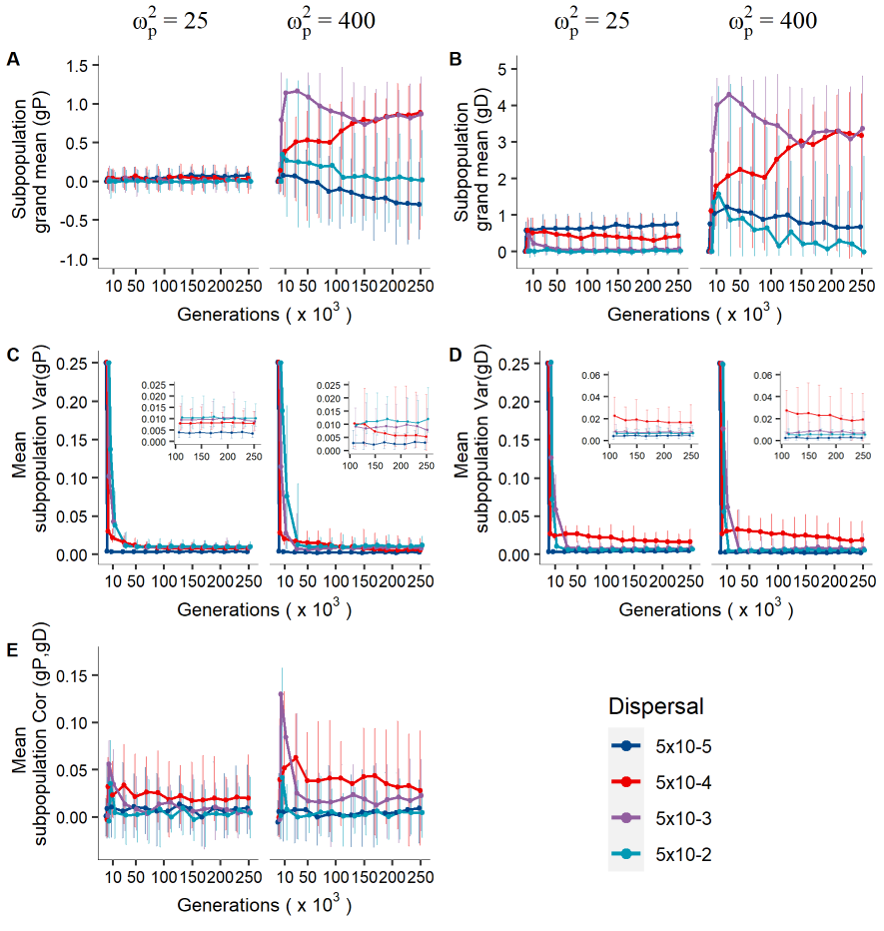

**Figure S6.** Effect of the strength of stabilising natural selection on female preference (ω^2^_P_ = 400, 25; weak and strong respectively) on the coevolution of costly female preference and male display under varying levels of dispersal. Panels, axes and lines as in Figure 1.

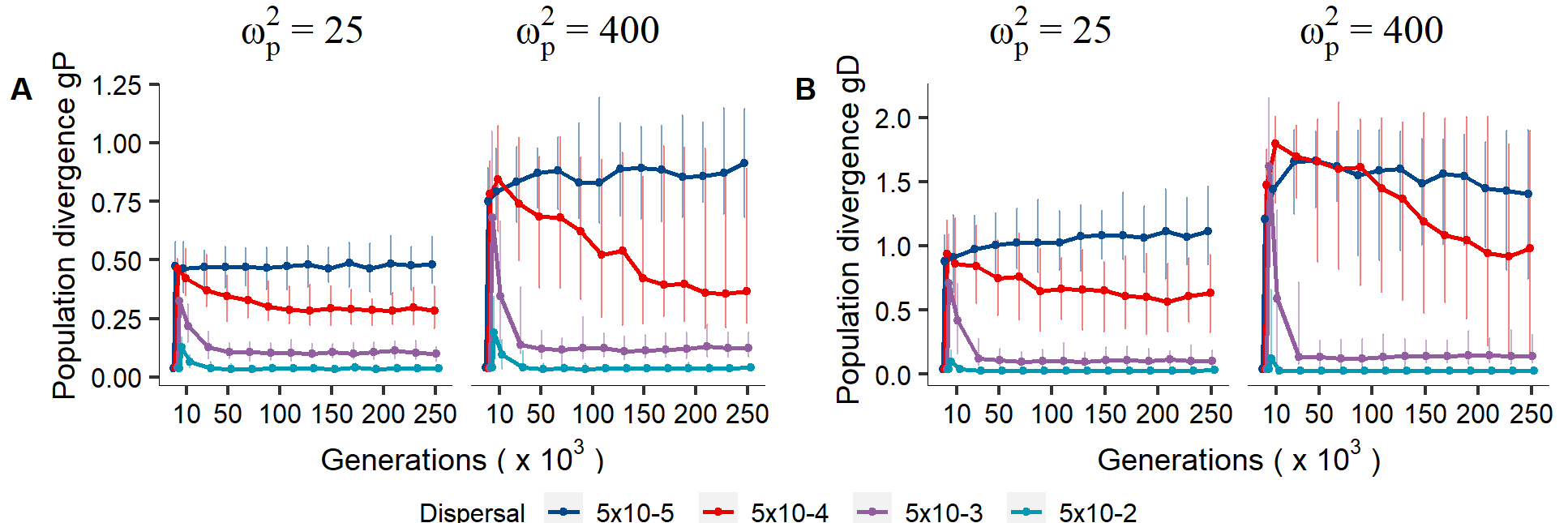

**Figure S7.** Effect of the strength of stabilising natural selection on female preference (ω^2^_P_ = 400, 25; weak and strong respectively) on population divergence in female preference under varying levels of dispersal. Panels, axes and lines as in Figure 3A and B.

### Effect of weaker stabilising natural selection on male display

We investigated the consequences of weaker stabilising natural selection on the male display trait on preference-display coevolutionary dynamics. Display generally evolved to higher values (Fig. S8B) but showed fluctuations through time under low dispersal. In accordance, preference shifted from positive to negative values for low levels of dispersal (Fig. S8A). Weaker direct selection against male display resulted in higher levels of population differentiation in display given low dispersal (*d* ≤ 5×10^-4^), while showing faster homogenisation given *d* > 5×10^-4^ (Fig. S8). However, reductions of subpopulation trait grand means and population divergence under low dispersal coincided with rapid shifts to population extinction (Fig. S25), suggesting that subpopulations with high female preferences and exaggerated display predominantly went extinct. This finding agrees with our main results showing that under high metapopulation subdivision and low dispersal, negative mean subpopulation preference coincided with high population extinction. Hence, extinction-recolonisation dynamics within metapopulations may cause fluctuations in the mean’s sexual traits that bear some similarity with previously described cycles of preference-display coevolution (Iwasa and Pomiankowski 1995).

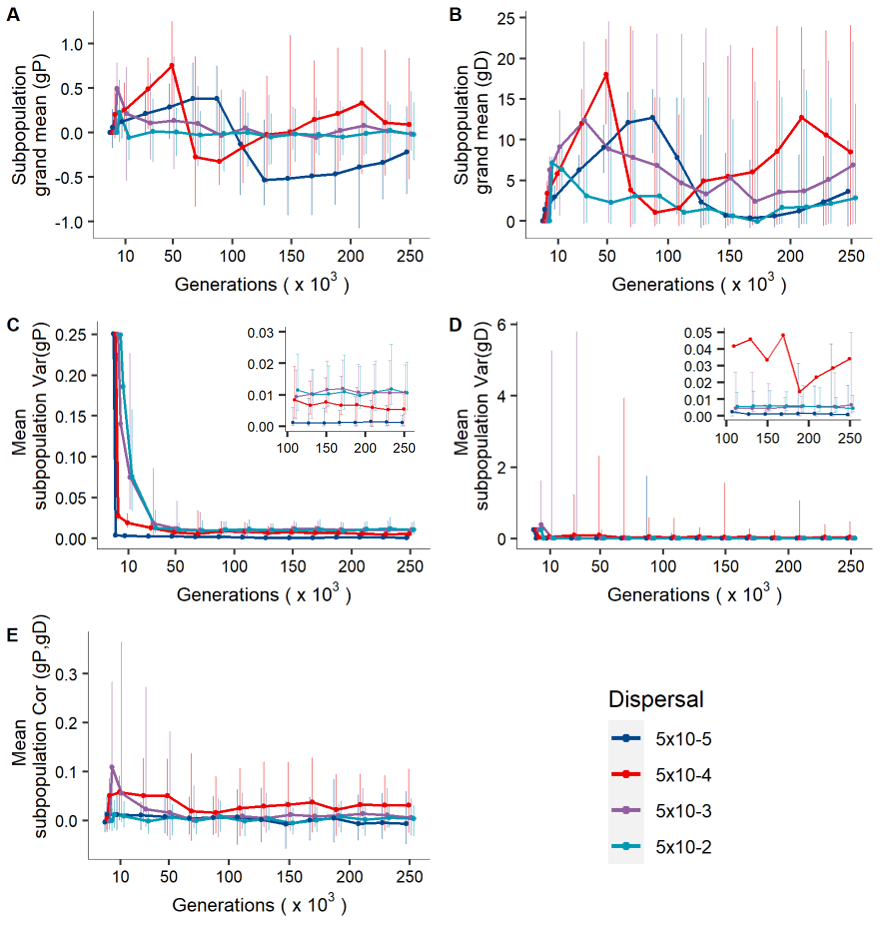

**Figure S8.** Effect of weaker stabilising natural selection on male display (ω^2^_D_ = 100) for coevolution of costly female preference and male display under varying levels of dispersal. Results are shown for metapopulations composed of 49 subpopulations over 250,000 generations. Panels, axes and lines as in Figure 1.

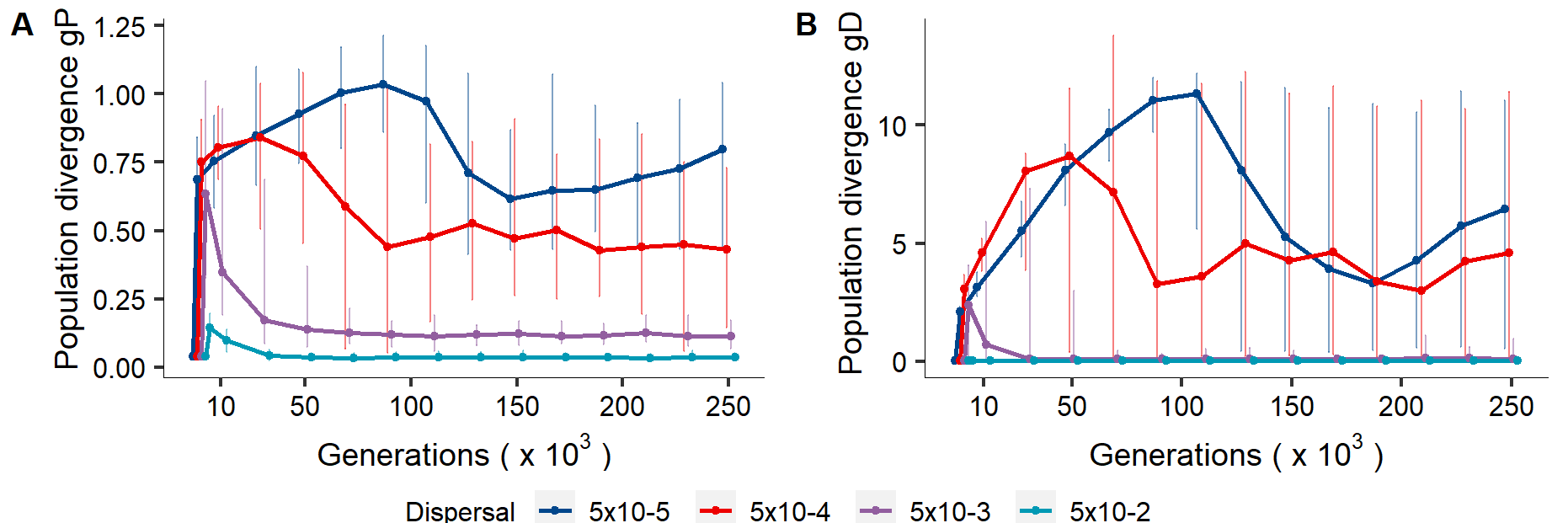

**Figure S9.** Population divergence in female preference genotype *gP* and male display genotype *gD* when stabilising natural selection on male display is relaxed (ω^2^_D_ = 100) in metapopulations composed of 49 subpopulations over 250,000 generations. Panels, axes and lines as in Figure 3A and B.

### Male optimal display phenotype θ_D_ = 10

Results remained qualitatively the same when we modelled a male display trait with the naturally selected optimum set to *θ_D_* = 10, allowing females with a negative mating preference to choose males of smaller than optimal phenotype. This led metapopulations to exhibit evolution towards either more positive or more negative preference and display (Fig. S10A-B), with a resulting grand mean of zero across replicate simulations. Genetic variation transiently increased in both sexual traits within metapopulations under dispersal *d =* 5×10^-2^ and was generally higher for display under *d =* 5×10^-4^ compared to our main analysis (Fig. S10D). The degree of population differentiation within metapopulations was also increased (Fig. S11).

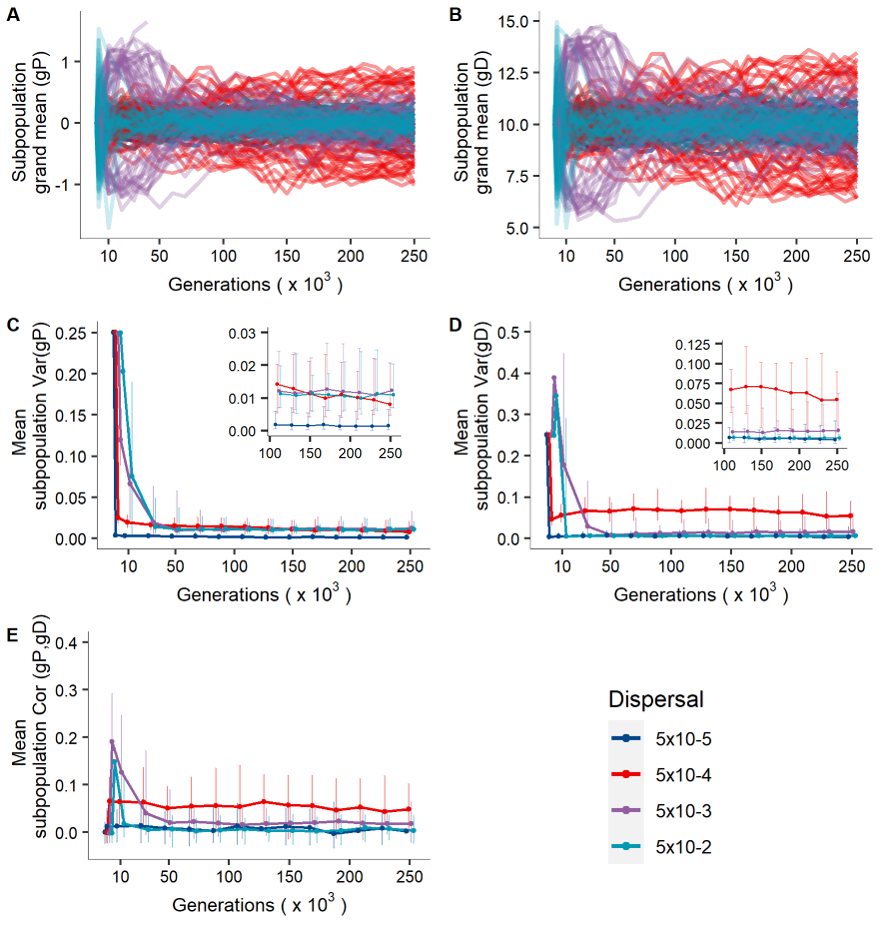

**Figure S10.** Coevolution of costly female preference and male display when both positive and negative preferences can evolve (i.e., male display naturally selected optimum *θ_D_* = 10). Results are shown for metapopulations composed of 49 subpopulations under varying levels of dispersal over 250,000 generations. In A-B, lines indicate single replicate simulations and are represented instead of the replicates grand mean which is zero given that single replicates can evolve either positive or negative female preference. Otherwise, panels, axes and lines as in Figure 1.

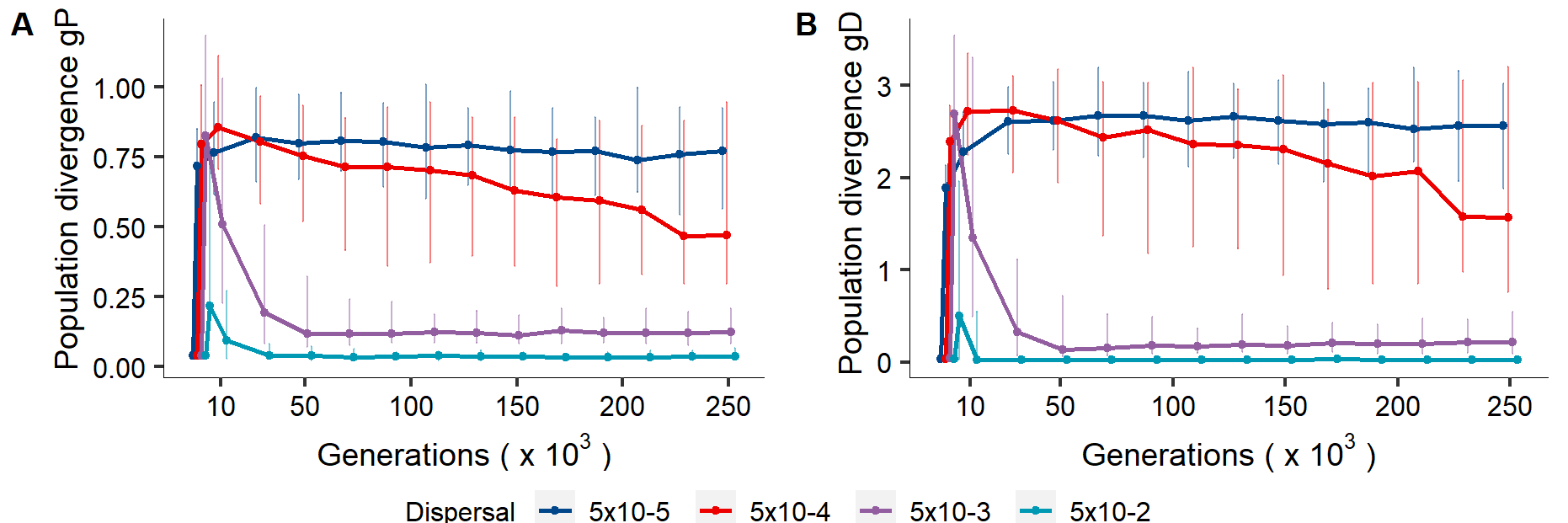

**Figure S11.** Population divergence in female preference genotype *gP* and male display genotype *gD* when both positive and negative preferences can evolve (i.e., male display naturally selected optimum *θ_D_* = 10). Results are shown for metapopulations composed of 49 subpopulations under varying levels of dispersal over 250,000 generations. Panels, axes and lines as in Figure 3A and B.

### Effect of the number of loci underlying each trait on preference and display coevolution

We tested the sensitivity of our model results to the number of loci, *L*, underlying each trait’s genotypic value. In addition to the results presented in the main text (*L* = 10), we simulated 100 loci. In our model, the mutational variance $V_{a_{M}}=2L\mu\sigma_{\alpha}^{2}$ (Walsh and Lynch 2019) is independent of *L*, due to the fact that we scale the variance of mutational effects ($\sigma_{\alpha}^{2}$) according to the number of loci $\sigma_{\alpha}^{2}= \frac{\sigma_{G,0}^{2}}{2L}*0.1$(Table S1). Results remained qualitatively similar with more loci underlying preference and display. The effect of more loci on the maintenance of V_a_ in display depended on dispersal probability. While V_a_ in display was lower under *d =* 5×10^-4^, V_a_ was marginally increased for *d =* 5×10^-3^ and *d =* 5×10^-2^ (Fig. S12 and Fig.S13).

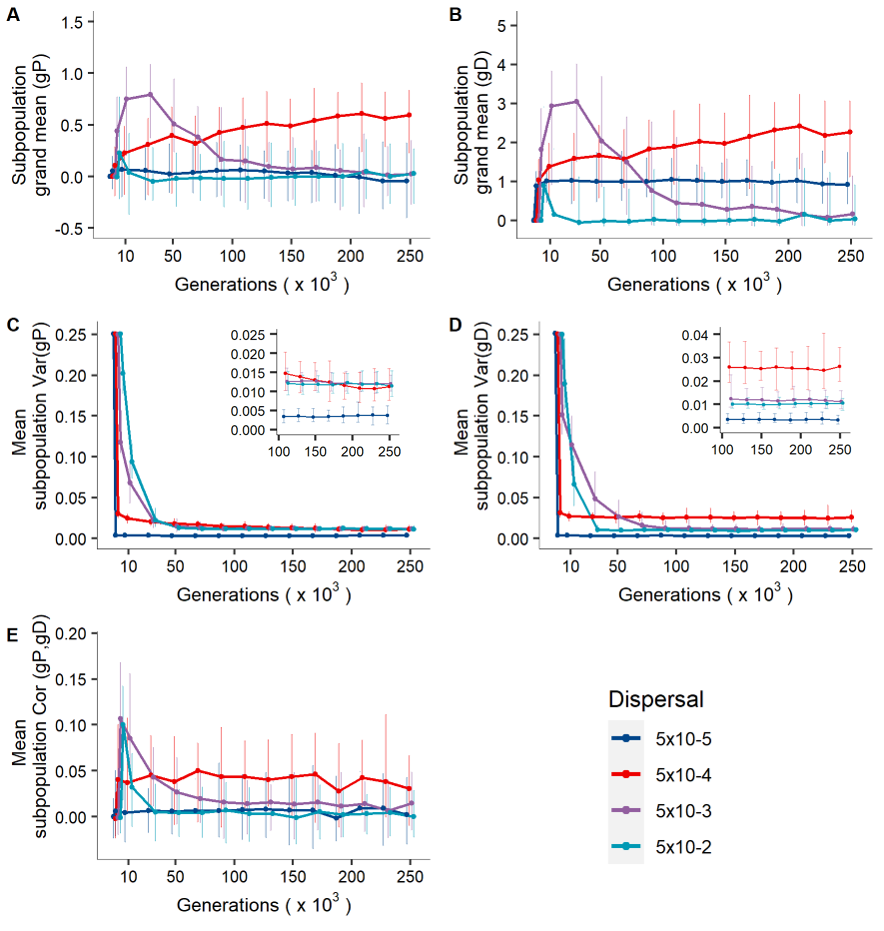

**Figure S12.** Coevolution of costly female preference and male display when the number of loci underlying the traits *L* = 100. Results are presented for metapopulations composed of 49 subpopulations over 250,000 generations. Panels, axes and lines as in Figure 1.

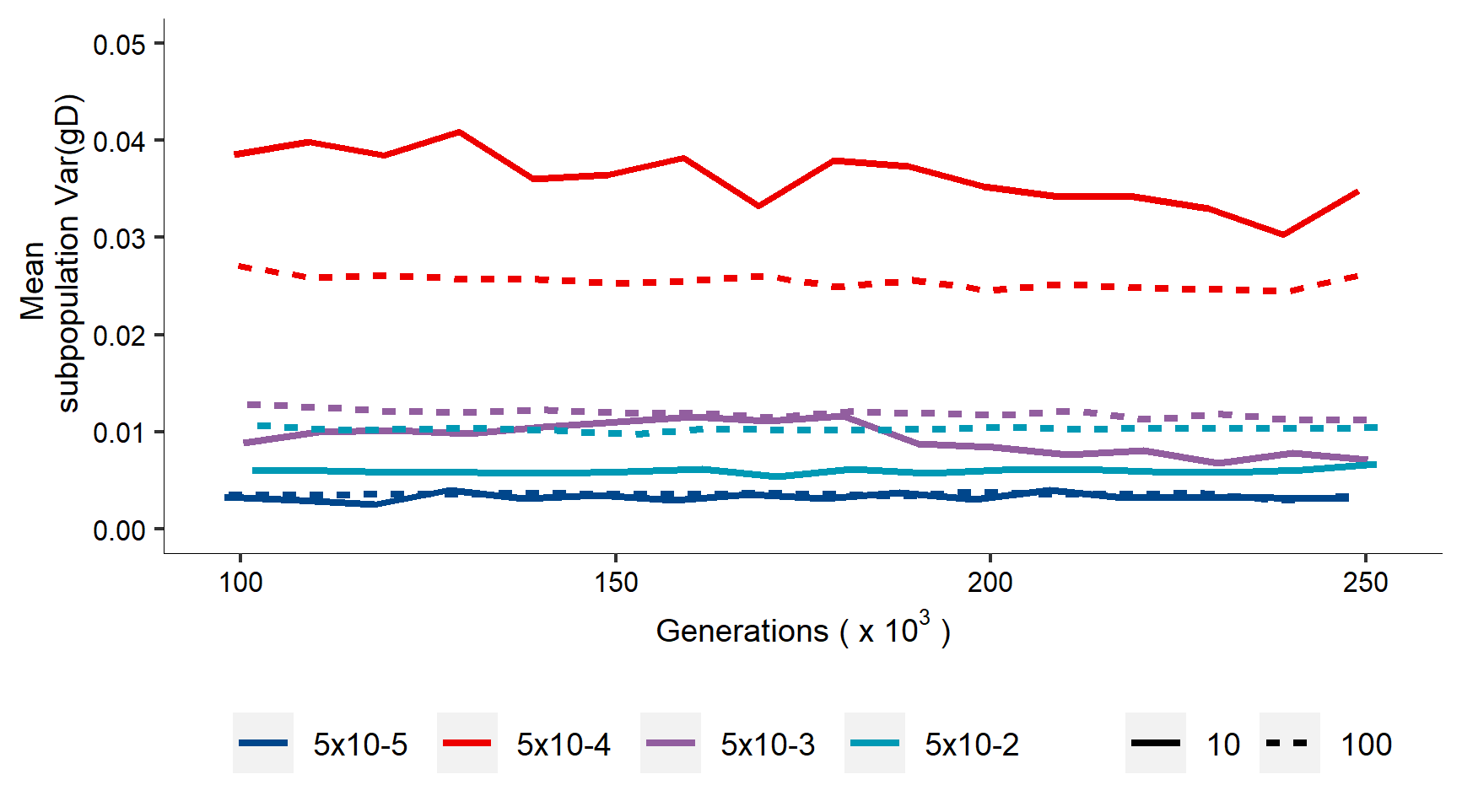

**Figure S13.** Comparison between *L* = 10 and *L* = 100 (solid and dashed line) on additive genetic variance in male display between generation 100,000 and 250,000. Lines indicate median across 50 replicate simulations.

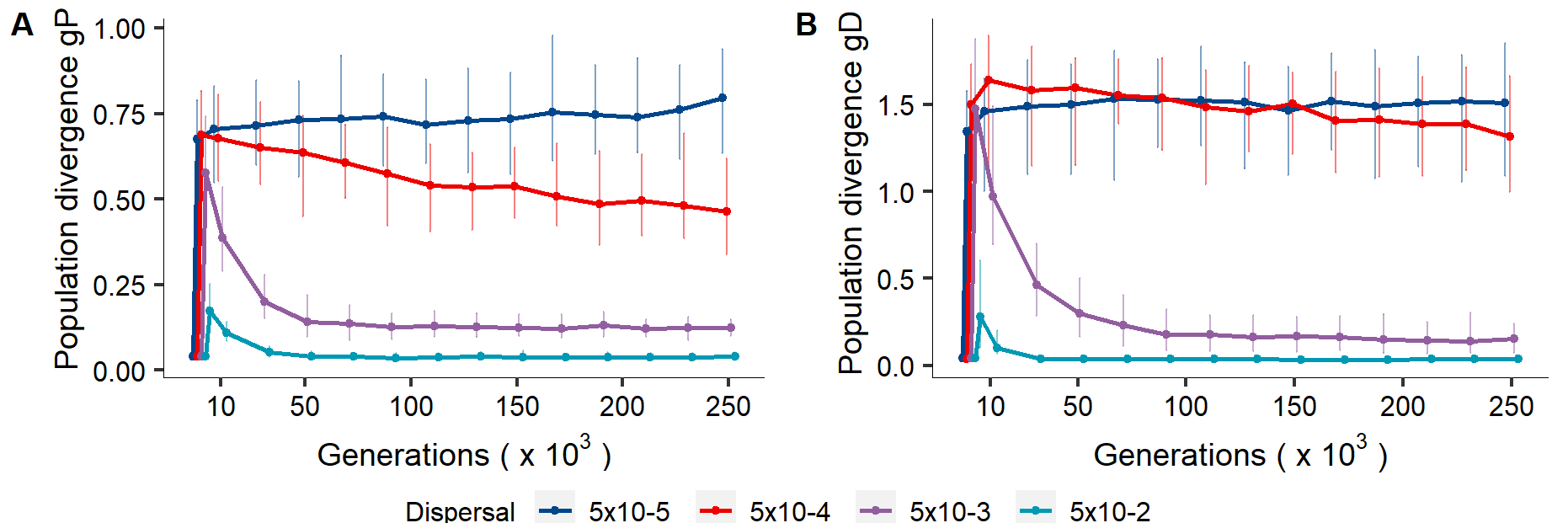

**Figure S14.** Population divergence in female preference genotype *gP* and male display genotype *gD* when number of loci underlying sexual traits is increased (*L* = 100) in metapopulations composed of 49 subpopulations over 250,000 generations. Panels, axes and lines as in Figure 3A and B.

### Preference and display coevolution under lower mutation probability

We further investigated how a hundredfold decrease in the probability of mutation would change our results (*μ =* 5×10^-6^/allele/generation). Transient dynamics were qualitatively similar to our main results. Female preference persisted over the 250,000 generations at both higher (*d* = 5×10^-3^) and lower (*d* = 5×10^-5^) dispersal. However, V_a_ in preference and display was eventually completely depleted under all dispersal scenarios.

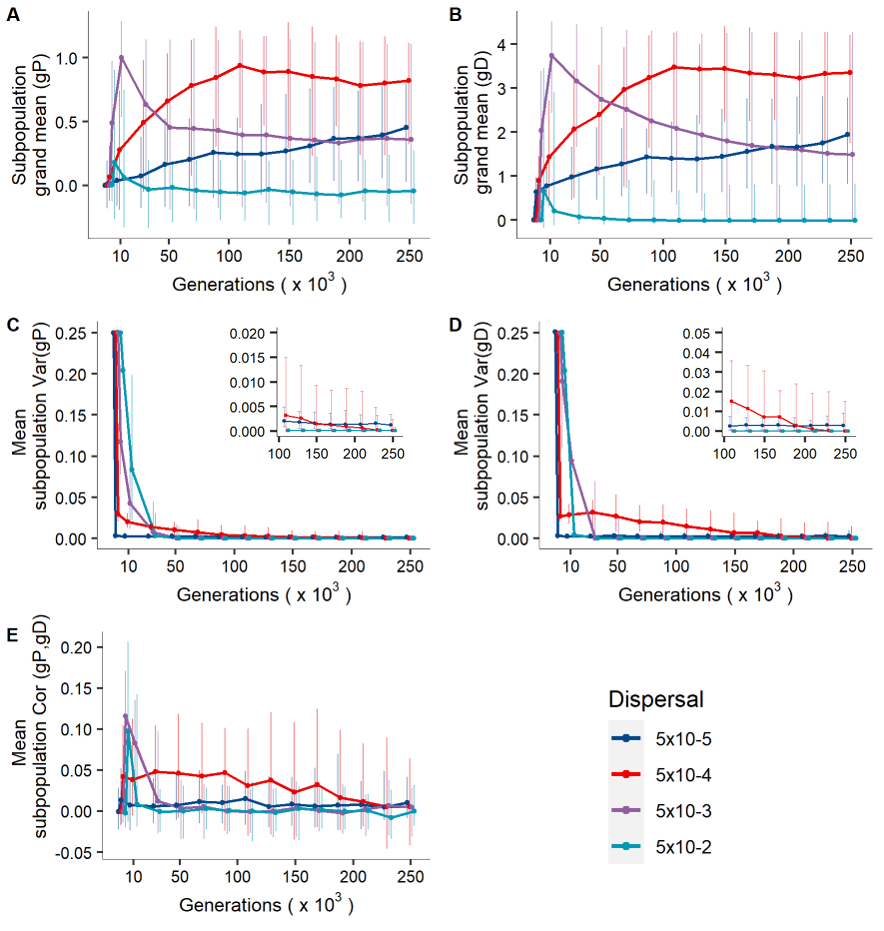

**Figure S15.** Effect of lower mutation rate (*μ* = 5×10^-6^/allele/generation) on the coevolution of costly female preference and male display under varying levels of dispersal. Results are shown for metapopulations composed of 49 subpopulations over 250,000 generations. Panels, axes and lines as in Figure 1.

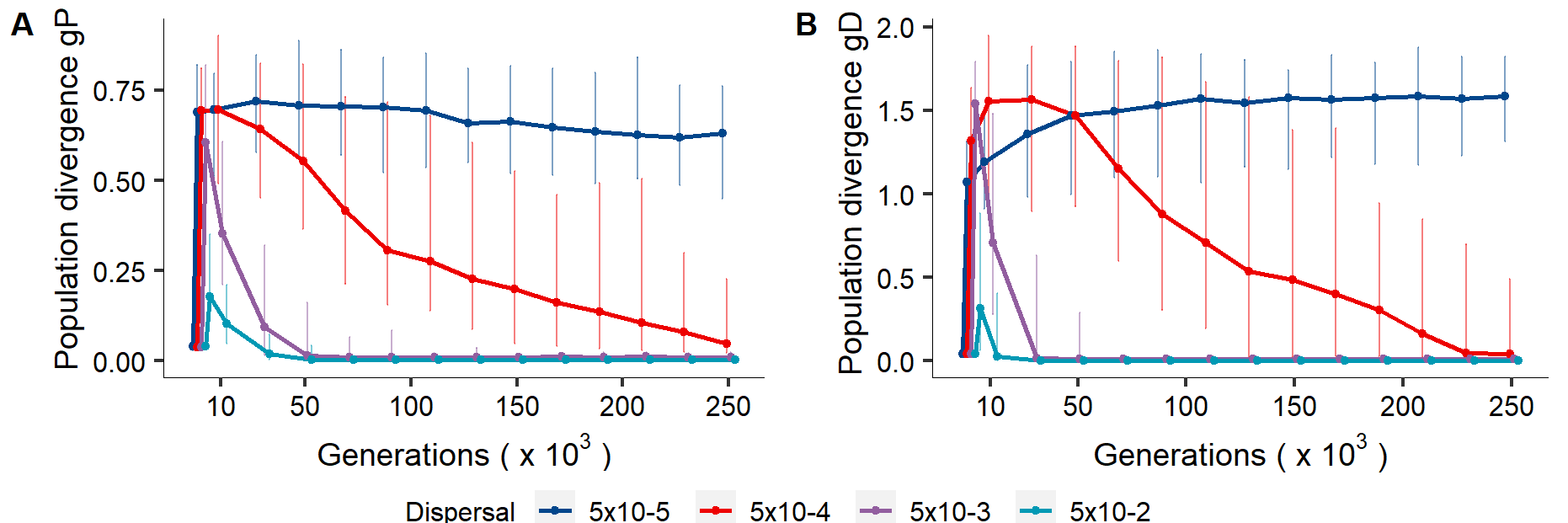

**Figure S16.** Population divergence in female preference genotype *gP* and male display genotype *gD* under mutation rate *μ=* 5×10^-6^/allele/generation in metapopulations composed of 49 subpopulations over 250,000 generations. Panels, axes and lines as in Figure 3A and B.

### Simulation starting with no standing genetic variation in preference and display

For simulations starting with some genetic variation in display and preference, we have shown that even under low mutational variance (the total amount of genetic variation arising each generation), variation in display persisted over long biological timeframes (Fig. S15). Here, we explore how much mutation is required to facilitate the evolution of costly preference and display by mutation accumulation alone, thus initialising preference and display at their naturally selected optima with no genetic variation. Simulations with mutation probability *μ* =5×10^-4^/allele/generation matched our main results (compare Fig. S17 with Fig. 1), indicating that the mutational variance is sufficient for the system to present similar short- and long-term dynamics. Simulations with *μ <* 5×10^-4^/allele/generation, without any initial input of standing genetic variation did not reach enough variation in display to promote evolution of costly preference. For simulations with *μ* =5×10^-4^, 5×10^-5^ and 5×10^-6^, mutational variance ($V_{a_{M}}=2L\mu\sigma_{\alpha}^{2}$) was 1.25×10^-5^ , 1.25×10^-6^ and 1.25×10^-7^ , respectively.

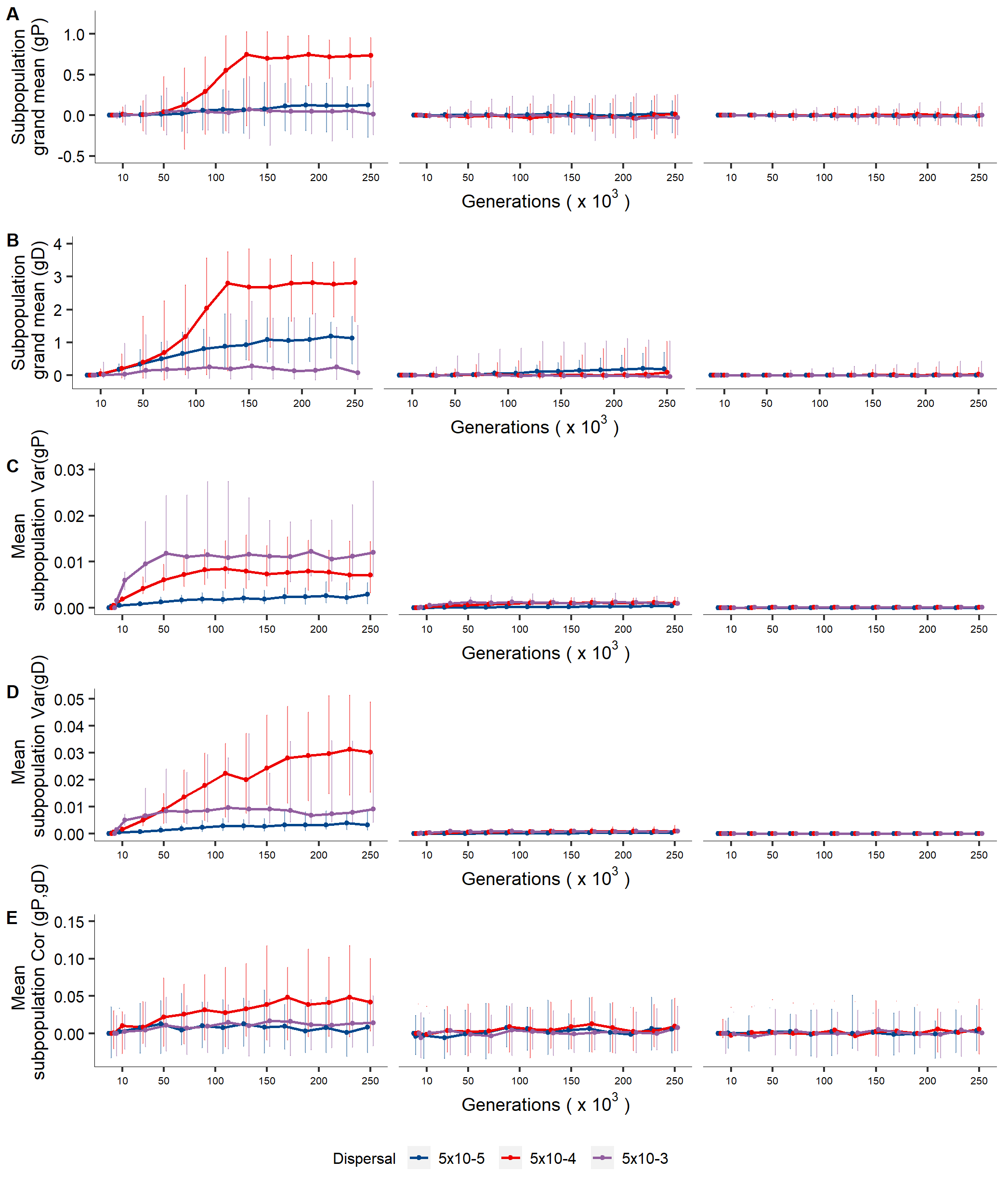

**Figure S17**. Coevolution of costly female preference and male display in the absence of initial genetic variation in preference and display. Results are shown for metapopulations composed of 49 subpopulations over 250,000 generations, under varying levels of dispersal and of mutation probability (left: *μ =* 5×10^-4^/allele/generation, middle *μ =* 5×10^-5^/allele/generation, right: *μ =*5×10^-6^/allele/generation). Panels, axes and lines as in Figure 1.

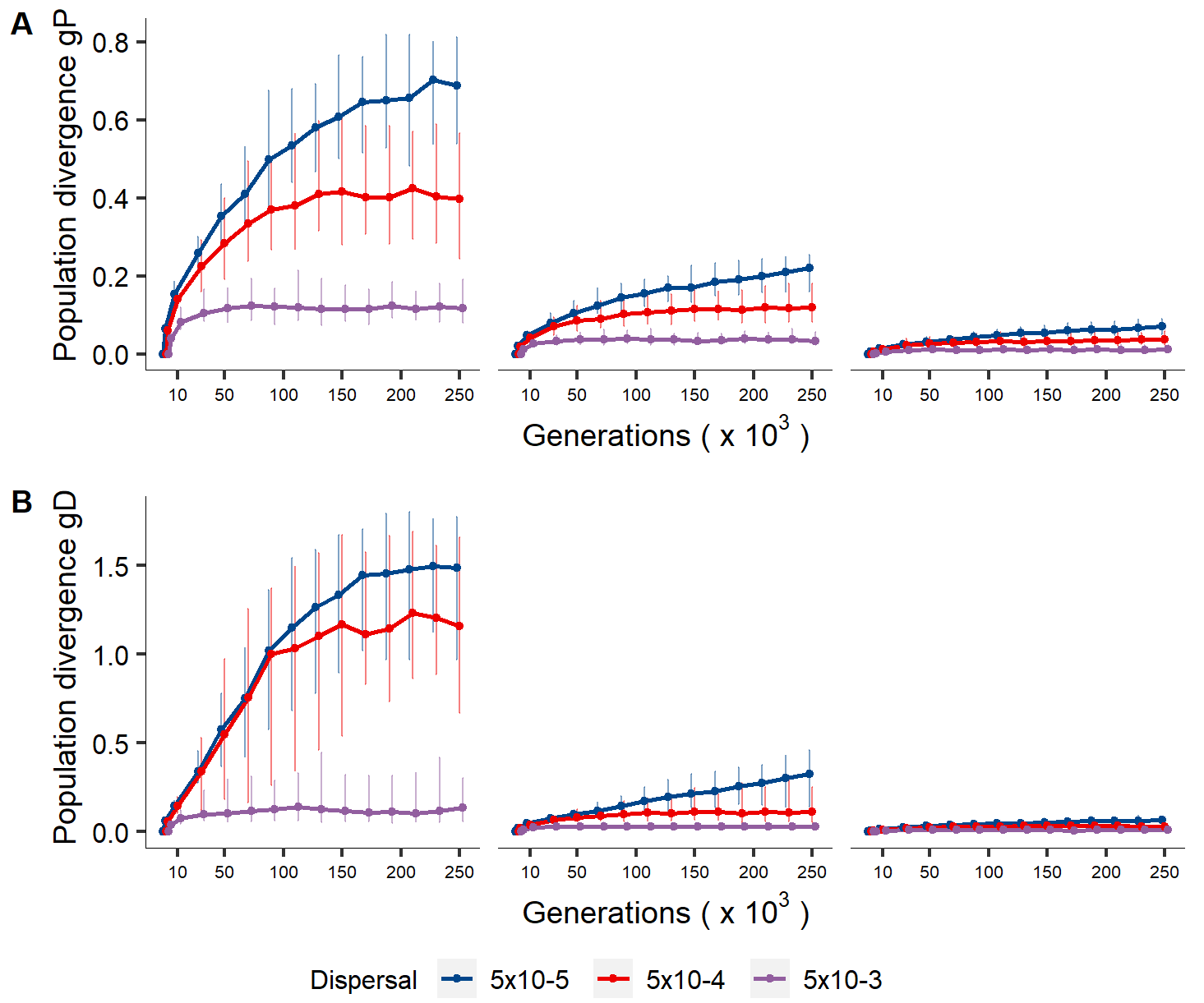

**Figure S18.** Population divergence in female preference genotype *gP* and male display genotype *gD* in metapopulations of 49 subpopulations over 250,000 generations. Simulations started without standing genetic variation for preference and display. Results are presented for three different values of mutation probability (left: *μ*=5×10^-4^/allele/generation, middle *μ*=5×10^-5^/allele/generation, right: *μ=*5×10^-6^/allele/generation). Panels, axes and lines as in Figure 3A and B.

### Evolutionary trajectories for different scenarios of metapopulation subdivision

Here we present the coevolutionary trajectories between female preference and male display for scenarios of metapopulation subdivision not shown in Figure 1 and Figure 3A-B of our main analysis.

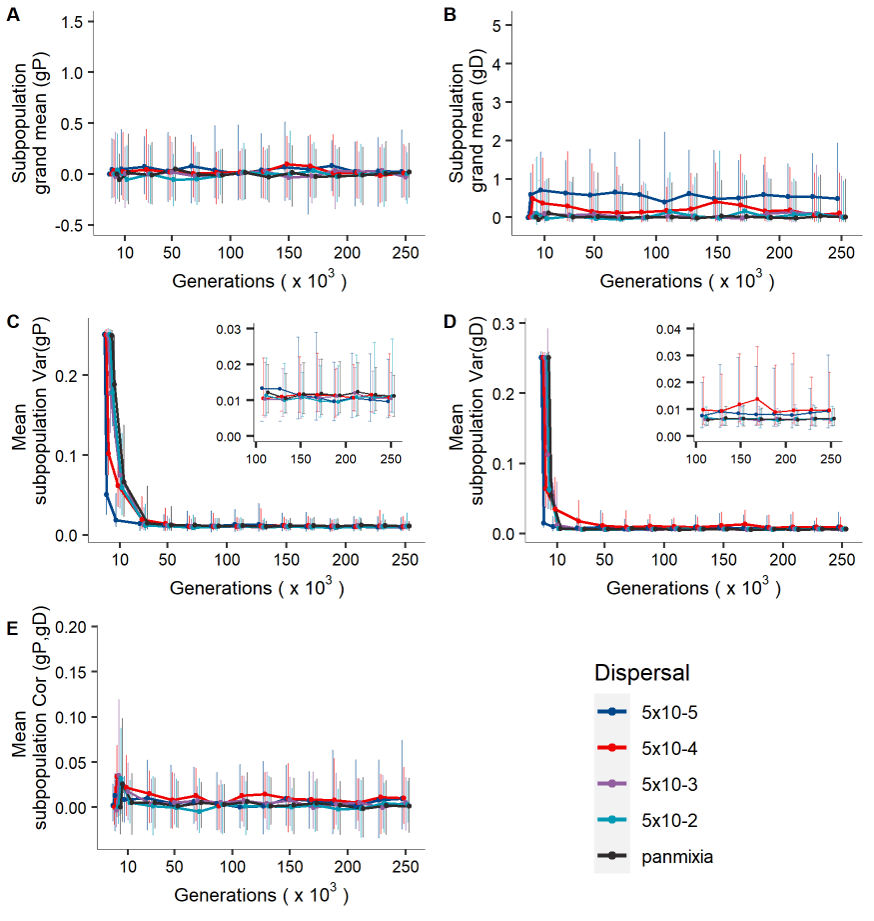
**Figure S19.** Coevolution of costly female preference and male display in metapopulations composed of **9 subpopulations** under varying levels of dispersal over 250,000 generations. Panels, axes and lines as in Figure 1.

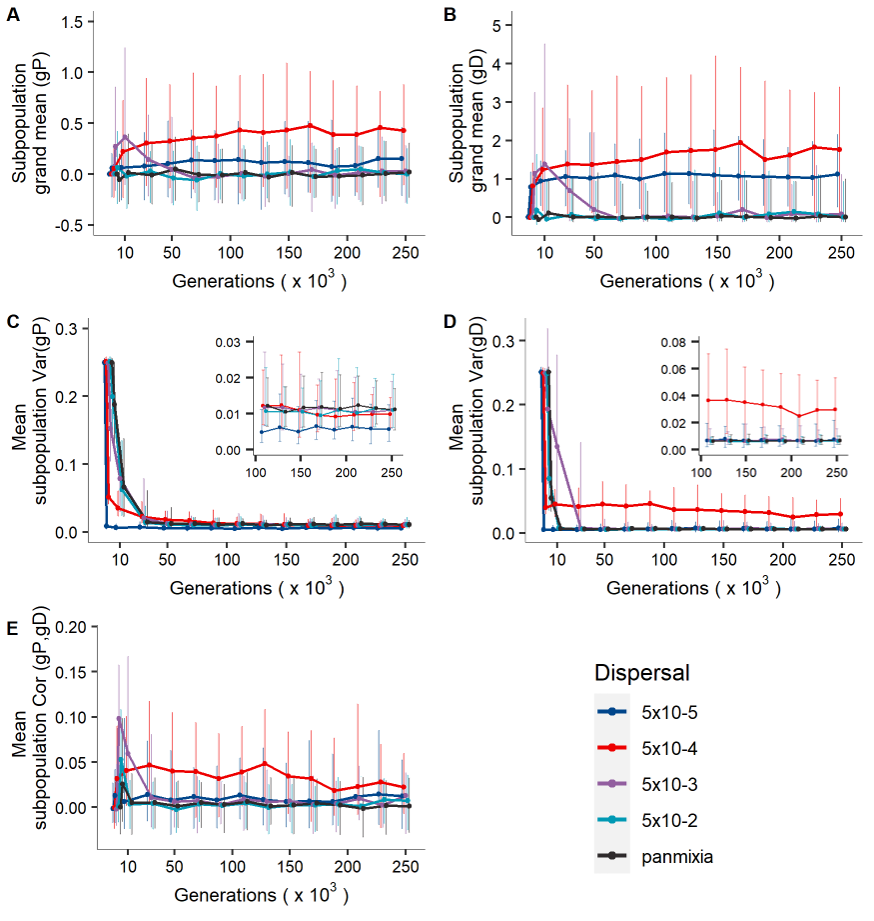

**Figure S20.** Coevolution of costly female preference and male display in metapopulations composed of **25 subpopulations** under varying levels of dispersal over 250,000 generations. Panels, axes and lines as in Figure 1.

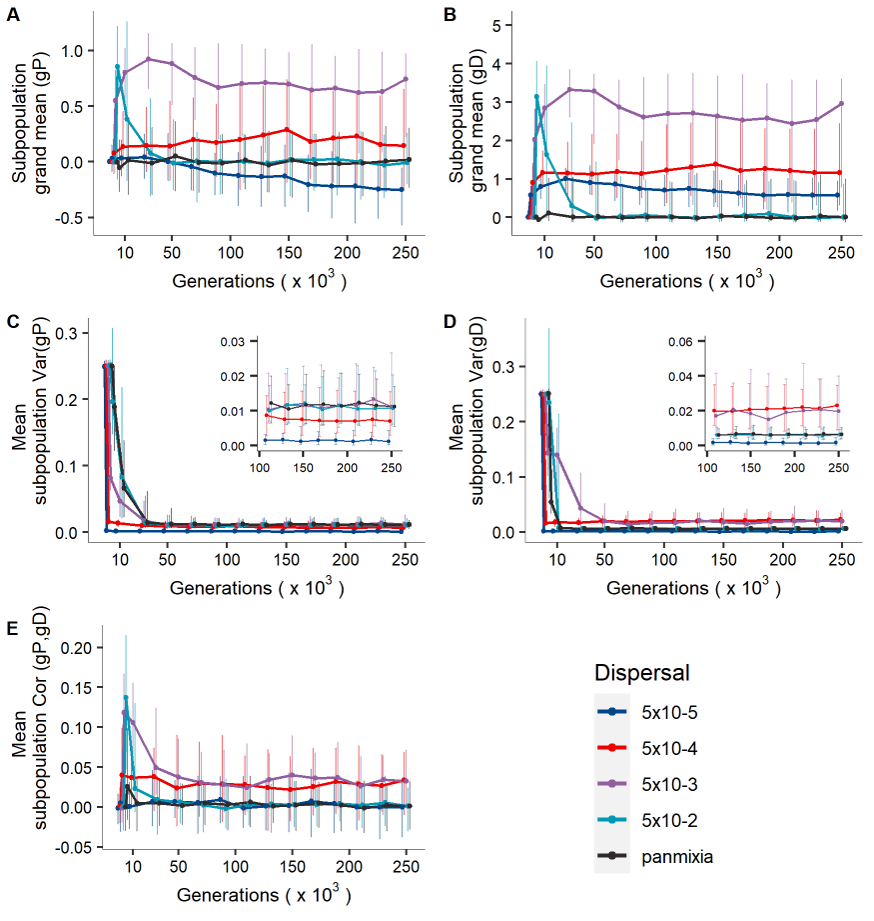

**Figure S21.** Coevolution of costly female preference and male display in metapopulations composed of **100 subpopulations** under varying levels of dispersal over 250,000 generations. Panels, axes and lines as in Figure 1.

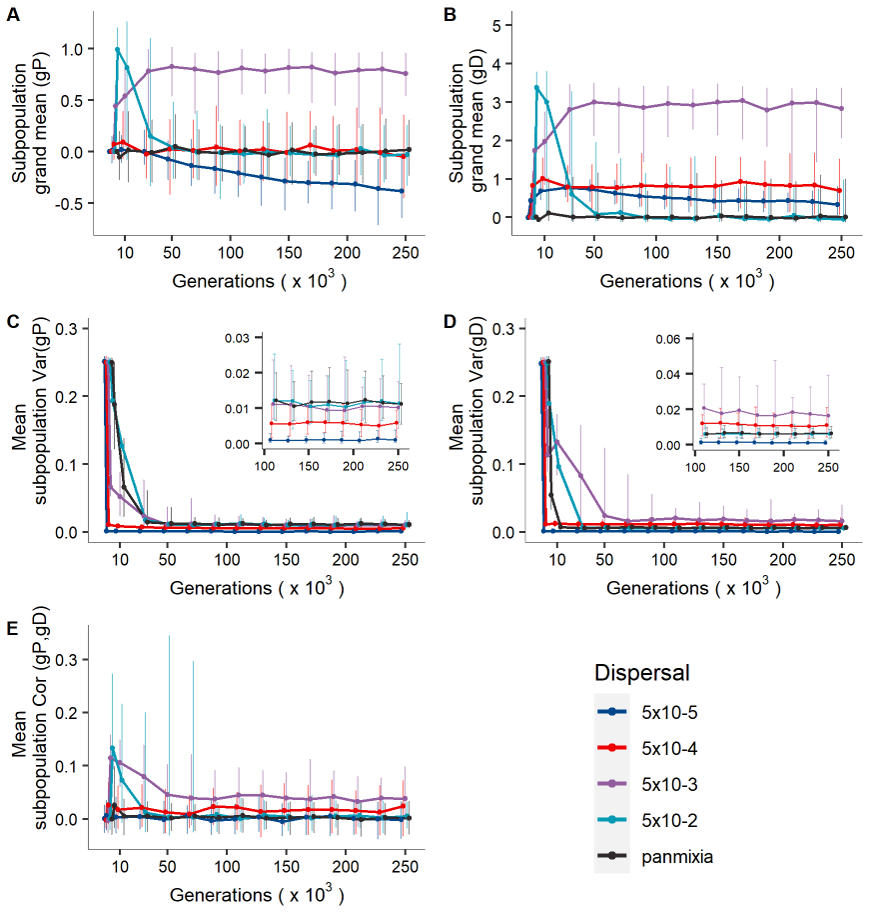

**Figure S22.** Coevolution of costly female preference and male display in metapopulations composed of **144 subpopulations** under varying levels of dispersal over 250,000 generations. Panels, axes and lines as in Figure 1.

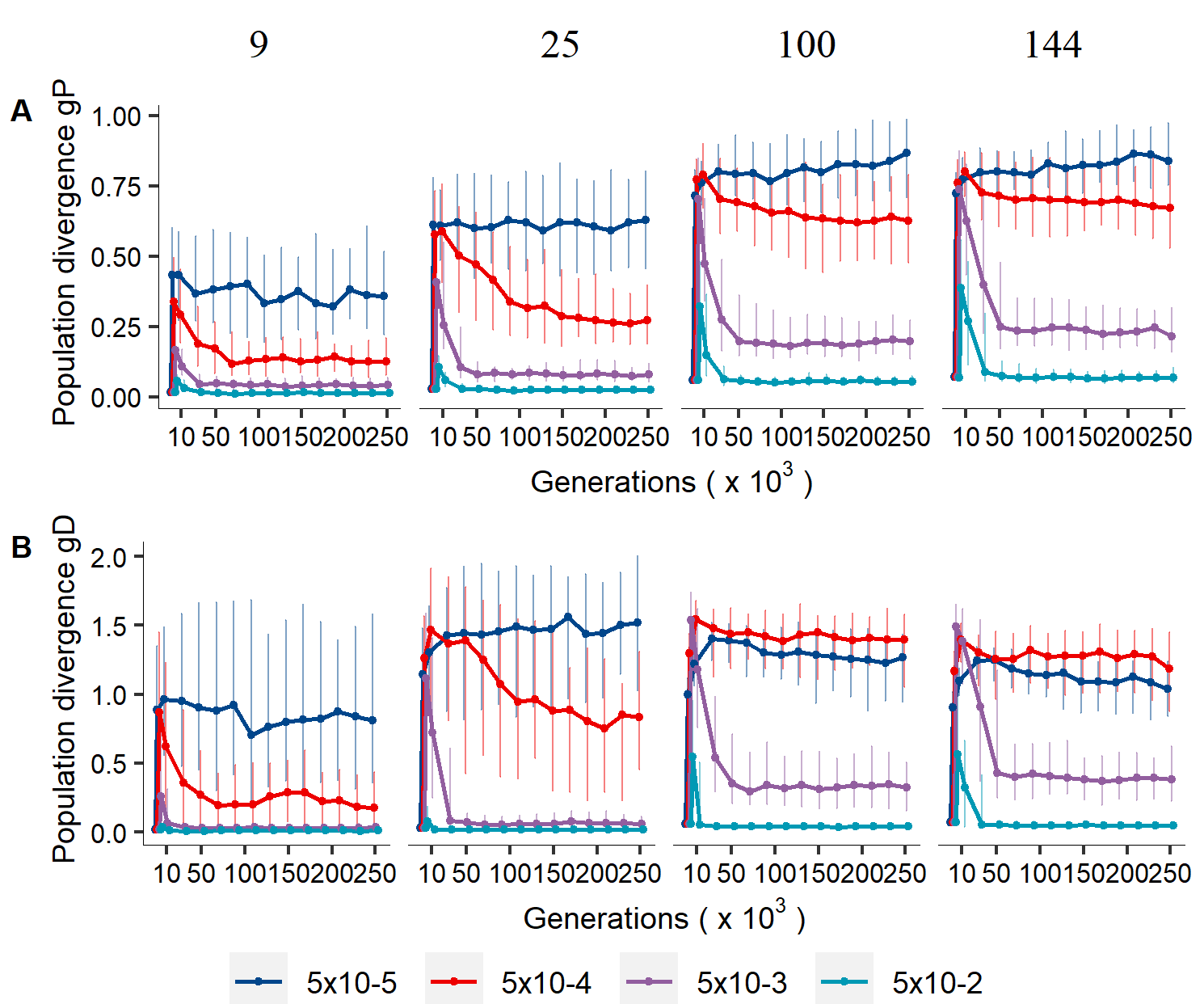

**Figure S23.** Population divergence in female preference genotype *gP* and male display genotype *gD* in metapopulations under varying levels of metapopulation subdivision (number of subpopulations 9, 25, 100, 144) over 250,000 generations. Panels, axes and lines as in Figure 3A-B.

### Extinction events due to sexual selection

In few cases, extinction of the whole metapopulation occurred (Fig. S24). Subpopulation extinction was most frequent under dispersal probability *d* = 5×10^-5^, while it was absent in simulation scenarios of random mating and in the panmictic population (not shown). Compared to a more gradual increase in subpopulation extinction in our main analysis (ω^2^_D_ = 4), relaxing the strength of natural selection on male display (ω^2^_D_ = 100) led, in some replicate metapopulations, to rapid extinctions after 50,000 to 100,000 generations (Fig. S25). Our results indicate that the nature of selection is crucial in determining the extinction-recolonisation dynamics. Different regimes of population regulation will likely also affect the source-sink dynamics within a metapopulation. Subpopulations closer to the naturally selected optimum may then more often act as source population, possibly even intensifying preference-display evolution in sink populations or completely swamping smaller subpopulations with low preference and display genotypes. The consequences for population extinction remain an exciting avenue for future research.

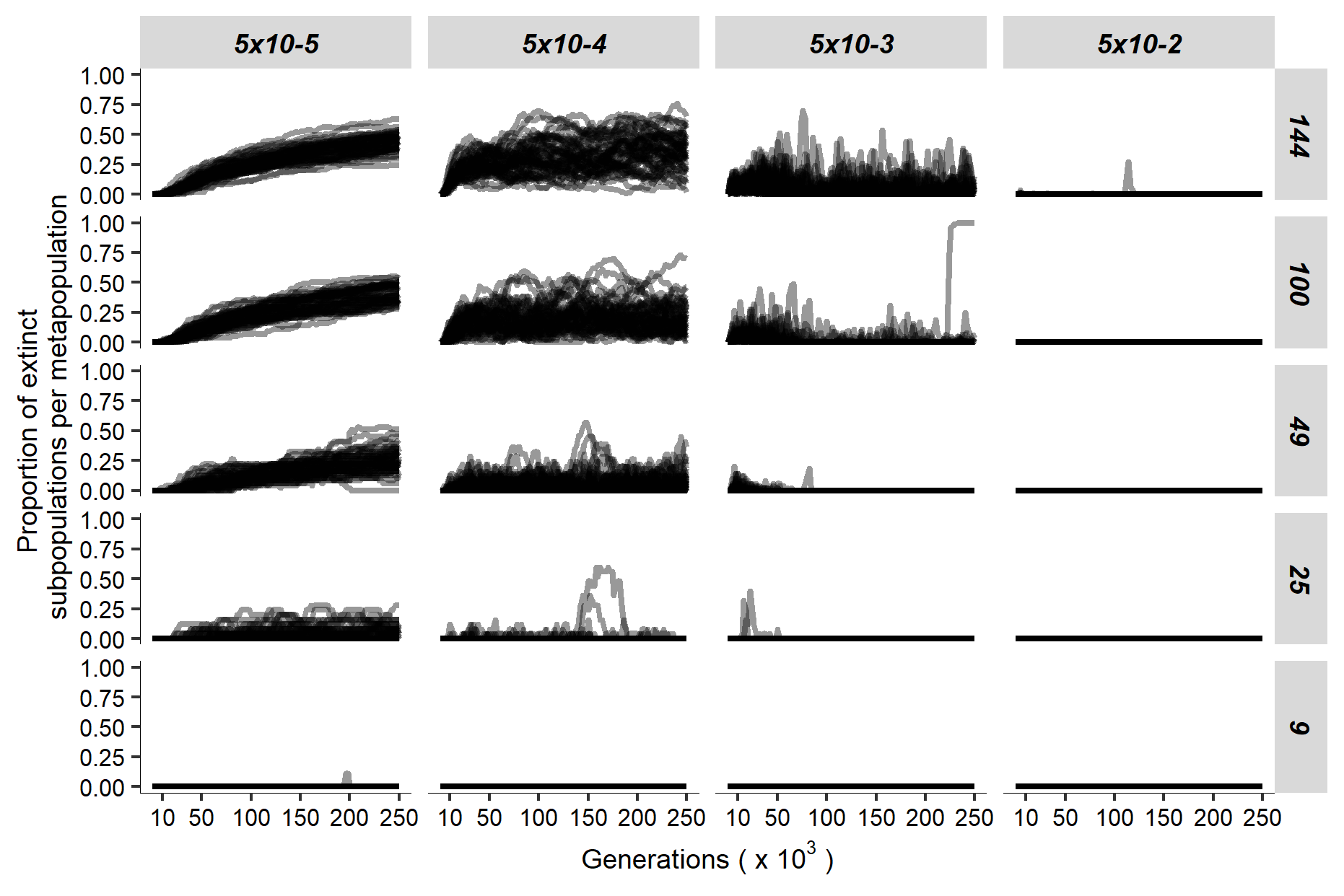

**Figure S24.** Proportion of subpopulations going extinct per metapopulation for different dispersal probabilities (columns) and across levels of spatial subdivision (rows). Lines indicate trajectories of single replicate simulations across 250,000 generations. Each simulation scenario was replicated 50 times and data is presented in intervals of 2,000 generations.

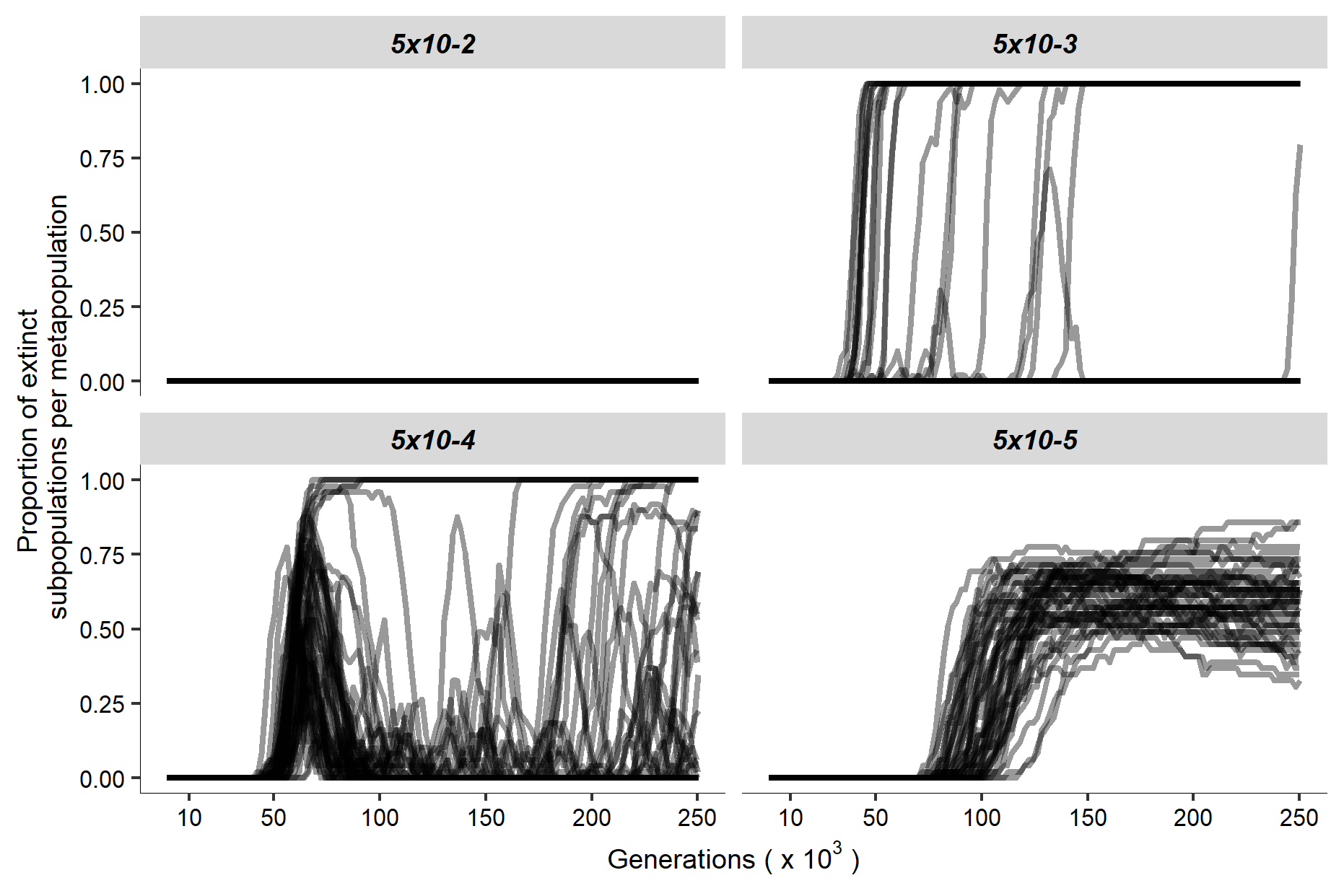

**Figure S25.** Proportion of subpopulations going extinct per metapopulation for different dispersal probabilities (different panels) in metapopulations composed of 49 subpopulations when stabilising natural selection on male display is relaxed (ω^2^_D_ = 100). Lines indicate trajectories of single replicate simulations across 250,000 generations. Each simulation scenario was replicated 50 times and data is presented in intervals of 2,000 generations.

### Biased dispersal associated with subpopulation display

We propose that extinction of populations exhibiting high preference-display genotypes could contribute to a migration bias by systematically underproducing emigrants, resulting in average emigrant display being lower than average subpopulation display. Indeed, within a metapopulation composed of 49 subpopulations, subpopulations exhibiting higher average display tend to produce less emigrants for *d* = 5×10^-4^ and 5×10^-3^ (Fig. S26B and C).

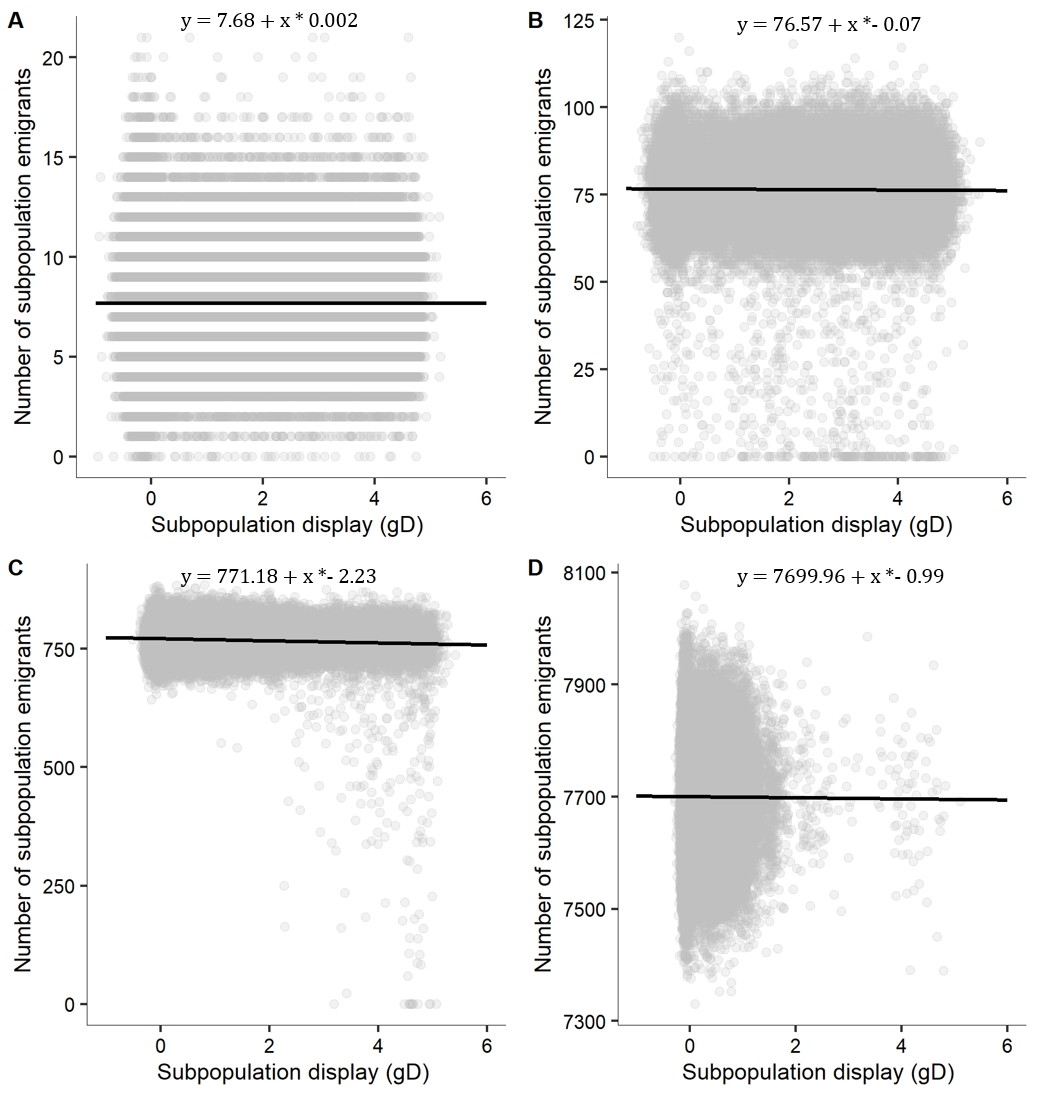

**Figure S26.** Absolute number of emigrants per subpopulation added-up over a time interval of 1,000 generations for different dispersal probabilities: A) *d* = 5×10^-5^; B) *d* = 5×10^-4^; C) *d* = 5×10^-3^ and D) *d* = 5×10^-2^. Line indicates linear regression of number of emigrants ~ display(gD). Points are extracted at generation intervals of 10,0000 across 250,000 generations from 50 replicate simulations.
